# Radiation and diversification of GATA-domain-containing proteins in the genus *Caenorhabditis*

**DOI:** 10.1101/2022.05.20.492891

**Authors:** Antonia C. Darragh, Scott A. Rifkin

## Abstract

Transcription factors are defined by their DNA-binding domains (DBDs). The binding affinities and specificities of a transcription factor to its DNA binding sites can be used by an organism to fine-tune gene regulation and so are targets for evolution. Here we investigate the evolution of GATA-type transcription factors (GATA factors) in the *Caenorhabditis* genus. Based upon comparisons of their DBDs, these proteins form 13 distinct groups. This protein family experienced a burst of gene duplication in several of these groups along two short branches in the species tree, giving rise to subclades with very distinct complements of GATA factors. By comparing extant gene structures, DBD sequences, genome locations, and selection pressures we reconstructed how these duplications occurred. Although the paralogs have diverged in various ways, the literature shows that at least eight of the DBD groups bind to similar G-A-T-A DNA sequences. Thus, despite gene duplications and divergence among DBD sequences, most *Caenorhabditis* GATA factors appear to have maintained similar binding preferences, which could create the opportunity for developmental system drift. We hypothesize that this limited divergence in binding specificities contributes to the apparent disconnect between the extensive genomic evolution that has occurred in this genus and the absence of significant anatomical changes.

## Introduction

Genomes are dynamic. Over time, processes like mutation, recombination, and gene conversion change genome sequences and can cause additions, losses, or relocations of genes within those genomes (Brown 2002). Over the last 50 years, many studies have demonstrated that gene duplication has served as a major mechanism through which new genes with novel functions evolve (Ohno 1970; Gottlieb 1977; Escriva et al. 2006; Assis and Bachtrog 2013; McKeown et al. 2014). For the first model of gene duplication, Ohno hypothesized that one of the duplicated gene paralogs would accumulate mutations while the other would maintain the original gene’s ancestral function since it would be constrained from accumulating mutations through negative selection (Ohno 1970). Because deleterious mutations are more frequent than beneficial ones (Charlesworth et al. 1994; Lynch and Walsh 1998; Eyre-Walker and Keightley 1999; Denver et al. 2004; Haag-Liautard et al. 2007) this scenario would likely result in the unconstrained paralog becoming non-functional (Nei and Roychoudhury 1973). However, if a beneficial mutation occurred and the new paralog gained a novel function, it could be preserved through positive selection. These two alternative outcomes for gene evolution are referred to as non-functionalization (or pseudogenization) and neofunctionalization, respectively (Ohno 1970).

Empirical evidence gathered since that first model of gene duplication was published (Ohno 1970) suggests that there are more paralogs per genome, but less pseudogenization, than Ohno’s theory predicted, (Allendorf, Fred W. et al. 1975; Ferris and Whitt 1979; Lundin 1993; Sidow 1996; Nadeau and Sankoff 1997; Postlethwait et al. 1998; Wendel 2000; Zhang 2003) and so a third outcome of gene duplication was proposed: subfunctionalization (Hughes 1994; Force et al. 1999; Lynch and Force 2000). Subfunctionalization is when duplicated genes each retain some, but not all, of the ancestral gene’s function such that negative selection preserves both genes in the genome. This process occurs through the accumulation of different and complementary deleterious mutations in both genes. If deleterious mutations occur more frequently than beneficial ones, subfunctionalization would be expected to occur more frequently than neofunctionalization and, depending on the specific function of the ancestral gene, at a similar or higher probability than would non-functionalization (Lynch and Force 2000). Another possible outcome of gene duplication is that the resulting gene paralogs may increase the organism’s robustness through biochemical redundancy or higher gene expression (Ohno 1970; Nei et al. 2000; Piontkivska et al. 2002; Kondrashov and Kondrashov 2006). In this scenario, both genes, each with the ancestral function, are retained in the genome. Teasing out paralog evolutionary histories is generally challenging since available information is often compatible with multiple possible histories.

The variability in the numbers of gene family members in different organisms is a testament to the pervasiveness and stochasticity of gene duplications (Ohno 1970; Jozefowicz et al. 2003; Baker and Woollard 2019). For example, diploid vertebrates have six to eight GATA-type transcription factors (GATA factors), arthropods and lophotrochozoans have four or five, and nematodes harbor from at least one to 36. Known nematode species with more than 14 genes encoding GATA factors are members of the genus *Caenorhabditis* (Lowry and Atchley 2000; He et al. 2007; Gillis et al. 2008; Tang et al. 2014; Eurmsirilerd and Maduro 2020; Maduro 2020). From a starting point of two GATA factors in the ancestor of Bilateria, two whole-genome duplications (Dehal and Boore 2005) and gene loss likely resulted in the six GATA factors currently found in mammals, and a third whole-genome duplication and gene loss likely led to the eight GATA factors found in teleost fish (Gillis et al. 2007; Gillis et al. 2008; Gillis et al. 2009). Evidence suggests that GATA factor evolution in arthropods and lophotrochozoans, which resulted in the four or five GATA factors found in contemporary species, occurred via tandem gene duplications (Gillis et al. 2008). Many tandem duplications of genes also likely occurred in nematodes since no evidence for whole-genome duplications has been found in any *Caenorhabditis* species to date (Semple and Wolfe 1999; Lynch and Conery 2000; Friedman and Hughes 2001; Cavalcanti et al. 2003; Stevens 2020). Changes in selective pressures likely preceded these duplications. There is evidence for positive selection on some sites in vertebrate GATA factors (Tang et al. 2014), and varying levels of selection on GATA factor DNA-binding domains (DBDs) have been proposed based on how conserved different residues are in these regions (Lowry and Atchley 2000; Maduro 2020). However, to our knowledge GATA factor families have not been tested for relaxed selection, which is expected under the subfunctionalization and pseudogenization gene duplication theories (Force et al. 1999) and which is common for recent paralogs (Lynch and Conery 2000).

The function of nematode GATA factors has mainly been studied in *Caenorhabditis elegans* (Eurmsirilerd and Maduro 2020). *C. elegans* GATA factors function in specific germ layers (Gillis et al. 2008), which is not necessarily the case for other GATA factors (Reuter 1994; Heitzler et al. 1996; Rehorn et al. 1996; Herranz and Morata 2001; Klinedinst and Bodmer 2003). *end-1*, *end-3, elt-7*, *elt-4*, and *elt-2* are exclusive to the endoderm, *med-1* and *med-2* function in both the endoderm and mesoderm, and *elt-1*, *elt-3*, *elt-6*, and *egl-18*, are all ectoderm-(hypoderm)-specific (Page et al. 1997; Zhu et al. 1997; Fukushige et al. 1998; Gilleard et al. 1999; Koh and Rothman 2001; Maduro et al. 2001; Maduro et al. 2005; McGhee et al. 2007). *elt-4*, a likely degenerate duplicate of *elt-2,* is expressed later in the development of the endoderm, but does not have any known function (Fukushige et al. 2003). We are publishing a detailed study on the evolution of the endoderm specification network elsewhere (Darragh AC, Rifkin SA, unpublished data, https://doi.org/10.1101/2022.05.20.492851, last accessed May 23, 2022). Here we present the evolution of the non-endoderm GATA factors (and other GATA-domain-containing proteins) with a focus on how their GATA DBDs have evolved.

Two studies of the evolution of GATA factors in nematodes, have been published recently. A comparison of GATA factor orthologs in 32 species of nematodes suggested that the genome of the ancestor of this phylum contained at least an *elt-1* ortholog and perhaps also an *elt-2* ortholog, and therefore that multiple gene duplications must have occurred since for evolution to have resulted in the 11 GATA factors currently encoded in the *C. elegans* genome (Eurmsirilerd and Maduro 2020). The other (Maduro 2020), like ours (Darragh AC, Rifkin SA, unpublished data, https://doi.org/10.1101/2022.05.20.492851, last accessed May 23, 2022) focused on the GATA factors that are expressed in the *C. elegans* endoderm. Dozens of new *Caenorhabditis* species have been discovered and sequenced over the last decade (Kiontke et al. 2011; Dey et al. 2012; Félix et al. 2014; Ferrari et al. 2017; Slos et al. 2017; Stevens et al. 2019; Stevens 2020; Teterina et al. 2020). Draft sequences of the genomes of an additional 34 *Caenorhabditis* species (Stevens 2020) are now available for carrying out even more comprehensive comparisons of GATA factors throughout the *Caenorhabditis* genus which should help elucidate the evolutionary history of this family of transcription factors.

Herein we report on our investigations into the evolutionary history of GATA-domain-containing proteins in 58 *Caenorhabditis* species and two outgroup nematode species. We tour the diversity and evolutionary history of GATA factors in the *Caenorhabditis* genus, highlighting selection pressures on paralogs. This study illustrates how closely related transcription factors have evolved and how gene duplications can be integrated into developmental regulatory networks, all without causing any obvious change in development or morphology.

## Results

### *Caenorhabditis* GATA-domain-containing proteins form twelve ortholog groups some of which cluster adjacently into larger clades

To identify potential GATA factors in *Caenorhabditis*, we searched for the characteristic GATA factor DBD motif defined by the PROSITE profile PS50114 (prosite.expasy.org; see Materials and Methods) in all fifty-six *Caenorhabditis* species for which genomic sequence assemblies were available, in *C.* sp. *45* and *C.* sp. *47* for which only transcriptome data were available, and in the genome assemblies of two outgroup *Diploscapter* species (*Caenorhabditis* Genome Project). We identified 890 protein-coding hits and made a preliminary estimation of their evolutionary relationships (data not shown). Because this PROSITE profile method unexpectedly left some species without GATA-domain-containing orthologs of *C. elegans* genes, we also used reciprocal protein BLAST tool (BLASTp) and translated nucleotide BLAST (tBLASTn), to search for missing orthologs by employing genes in their sister species as bait (Altschul et al. 1990; Camacho et al. 2009). This reciprocal BLAST analysis and edits of errors that we identified in multiple gene annotations (see Materials and Methods), identified 51 additional proteins which brought our total up to 941 genes (Supp. Table 1). We estimated the evolutionary history of the 884 well-alignable (see Materials and Methods) proteins using maximum likelihood approaches (Kalyaanamoorthy et al. 2017; Minh et al. 2020) and a GATA factor from the slime mold *Dictyostelium fasciculatum* to root our phylogenetic analysis (Fig. 1). Some GATA-domain-containing proteins are highly divergent, with long branches that do not group robustly into any clade in our phylogeny (Fig. 1). Excluding these and other probable pseudogenes (Supp. Fig 1; see Materials and Methods) left us with 714 GATA-domain-containing protein sequences in which we had high confidence (Supp. Table 1). We focused on these proteins for further analysis. We also re-estimated the evolutionary history of this subset of proteins, which produced a similar topology and will be reported elsewhere.

**Figure 1.**
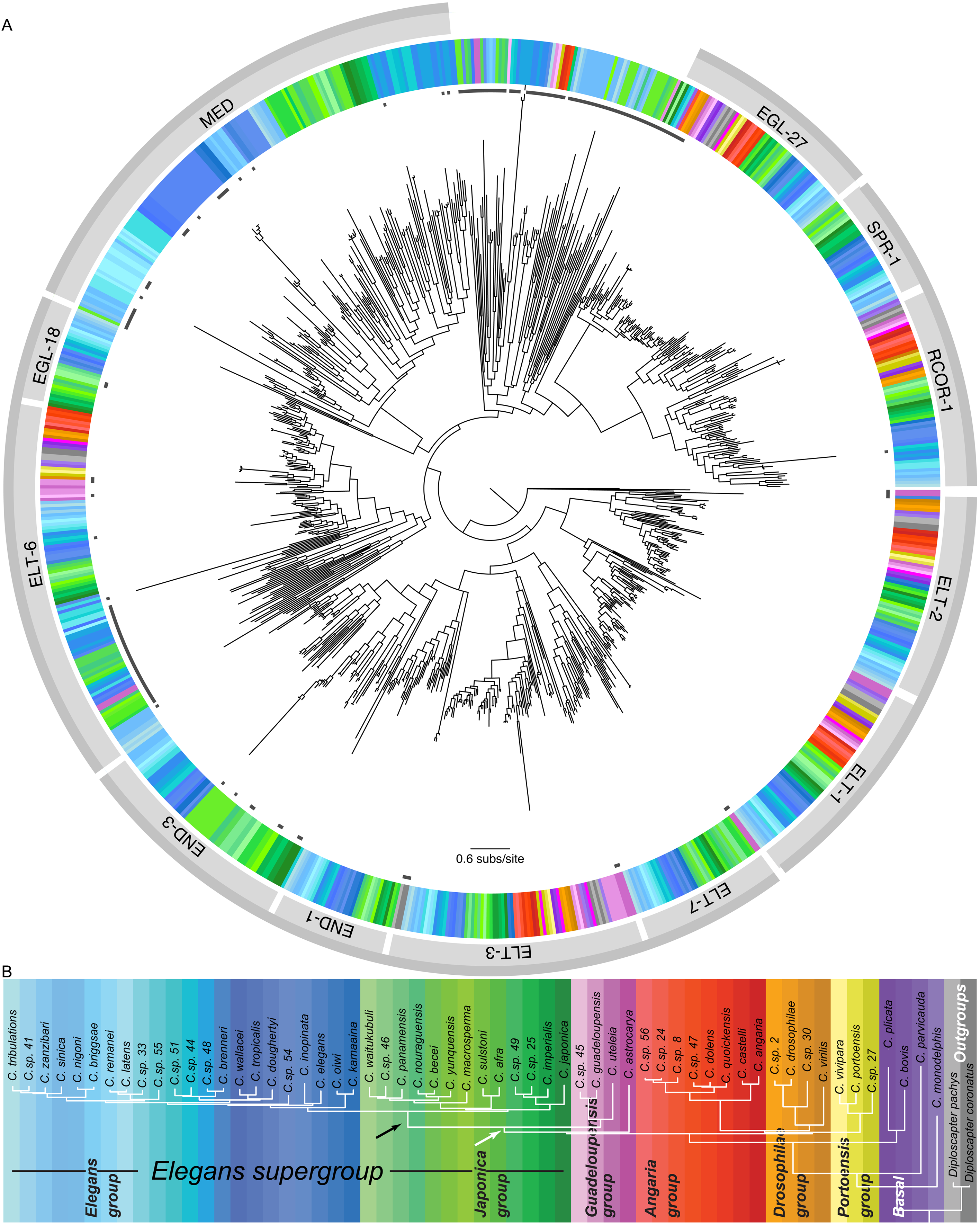
Preliminary inferred evolutionary history of *Caenorhabditis* GATA-domain-containing proteins. **(A)** Maximum likelihood phylogeny of 884 alignable GATA-domain-containing proteins in 58 *Caenorhabditis* and two outgroup nematode species, created using a GATA factor from the slime mold *Dictyostelium fasciculatum* to root the tree. The colors in the ring encircling the tree correspond to the species in which the proteins in the tree were identified; the key to color-species correspondence is given in B below. The names of the 12 ortholog groups the 884 proteins were categorized into (see Results) are indicated in the lighter of the two outer gray rings (with white gaps between groups). Groupings of ortholog groups that share adjacent clades in the tree are highlighted by a darker gray line on the outermost edge of the figure (with white gaps between groups). The broken ring of small black rectangles (under the species color ring) indicate GATA factors in which we had low confidence and were excluded from further analyses. The key for translating branch length into evolutionary distance (in units of amino acid substitutions per site) is shown near the bottom of the phylogenetic tree. **(B)** Phylogenetic relationships among the 60 species used in this study (based on Stevens 2020). Each species is designated by a different color shade; the same color-species designations are used in A above. The black arrow points to the *Elegans* supergroup ancestral branch where the ancestral *spr-1* gene likely arose. The white arrow points to the *Elegans* supergroup/*Guadeloupensis* group ancestral branch where the ancestral *egl-18* gene likely arose.

Our resulting phylogenetic tree contains 12 distinct groups of orthologs, each group named after the *C. elegans* protein(s) within that group (Fig. 1A). The clade designations in our tree are similar to those established for *Caenorhabditis* GATA factors previously (Gillis et al. 2008; Eurmsirilerd and Maduro 2020; Maduro 2020). However, because we have included many more *Caenorhabditis* species, all known *Caenorhabditis* GATA factors, and additional *Caenorhabditis* proteins that contain GATA-like domains (e.g., EGL-27, RCOR-1, and SPR-1 orthologs), our tree represents a more complete picture of how *Caenorhabditis* GATA factors evolved. The clades in our phylogenetic tree are robust against ultrafast bootstrapping (Minh et al. 2013; Hoang et al. 2018) and have well-supported nodes (Fig. 1A). A striking feature of our phylogeny is that GATA-domain-containing proteins have radiated extensively within the *Elegans* supergroup, as exemplified by the END-3, END-1, ELT-7, MED, and SPR-1 orthologs, which are unique to the *Elegans* supergroup and by EGL-18 which is also found in the *Guadeloupensis* group (Fig. 1A). This pattern for END-3, END-1, ELT-7, and MED orthologs was first identified by Maduro (2020) and then supported by the lack of orthologs for these proteins in other nematodes (Eurmsirilerd and Maduro 2020). We extend this finding to 34 more *Caenorhabditis* species.

Ten of the 12 ortholog groups clustered into four larger clades of adjacent ortholog groups (Fig. 1A). We refer to these larger clades by the name of the most ancient ortholog group within the clade or by the names of all the ortholog groups included in the clade when the most ancient group is not apparent. There are four of these larger clades. These clades include the rcor1 clade which contains the RCOR-1 and SPR-1 ortholog groups, the elt6 clade which contains the EGL-18 and ELT-6 ortholog groups, the elt1/2 clade which contains the ELT-1 and ELT-2 ortholog groups, and the elt3 clade which is composed of the ELT-3, ELT-7, END-1, and END-3 ortholog groups (Fig. 1A). We present below detailed results for the non-elt3 clades and for the EGL-27 ortholog group and include the elt3 clade and MED ortholog group for a comparison of all *Caenorhabditis* GATA DBDs. More details of the elt3 clade and MED ortholog group evolution will be reported elsewhere (Darragh AC, Rifkin SA, unpublished data, https://doi.org/10.1101/2022.05.20.492851, last accessed May 23, 2022).

### Protein groups with non-canonical GATA domains and non-GATA domains

#### rcor1 clade

##### An rcor-1 gene duplication in the *Elegans* supergroup ancestor likely produced the ancestral *spr-1* gene

The SPR-1 and RCOR-1 ortholog groups form a well-supported (100% ultrafast bootstrap support (Minh et al. 2013; Hoang et al. 2018)) monophyletic clade in our phylogenetic tree (Fig. 1A), which supports their shared homology to the same proteins in other nematodes, CoRest-PI in *Drosophila melanogaster*, and components of the REST corepressor complex in vertebrates (Wormbase.org). In *C. elegans*, SPR-1 and RCOR-1 play non-redundant roles in differentiation of pi cells during gonad and vulval development (Jarriault and Greenwald 2002; Bender et al. 2007; Hale et al. 2014; Vandamme et al. 2015). We only found SPR-1 orthologs in *Elegans* supergroup species, whereas RCOR-1 orthologs were identified in species throughout the *Caenorhabditis* genus as well as in the two *Diploscapter* species included in our analysis (Fig. 1). These results (and others described below) support the hypothesis that a *rcor-1* duplication produced *spr-1* in the ancestor of the *Elegans* supergroup.

##### *spr-1* and *rcor-1* orthologs have similar gene structures including non-GATA domains

We compared the structures of extant *spr-1* and *rcor-1* genes (Supp. Fig. 2A,B) and used parsimony to predict the *Elegans* supergroup *rcor-1* and *spr-1* ancestral structures (Fig. 2A). These predicted structures are very similar, the main difference being that the *spr-1* exon 9 is about three times longer than that of *rcor-1* (Fig. 2A). In fact, this exon, or an exon around this position, is more variable in length in extant *spr-1* genes, leading to more variability in the location of the second Myb/SANT (Swi3/Ada2/N-CoR/TFIIIB) (Aasland et al. 1996) domain compared to *rcor-1* genes (Supp. Fig. 2A,B). Two distinguishing features of most *spr-1* and *rcor-1* genes is that they have a conserved splice site immediately following the first 36 nucleotides that code for their atypical GATA ZnFs (see below) and a conserved splice site after the first 84 nucleotides coding for their first Myb/SANT domain (Fig. 2A; Supp. Fig. 2A,B). These conserved splice sites are not found in any of the other GATA-domain-containing proteins included in this study (Supp. Fig. 2A,B), thus providing further support for the common ancestry between *spr-1* and *rcor-1* genes and for the distant placement of the rcor1 clade away from canonical GATA factor groups in our phylogeny (Fig. 1A).

**Figure 2.**
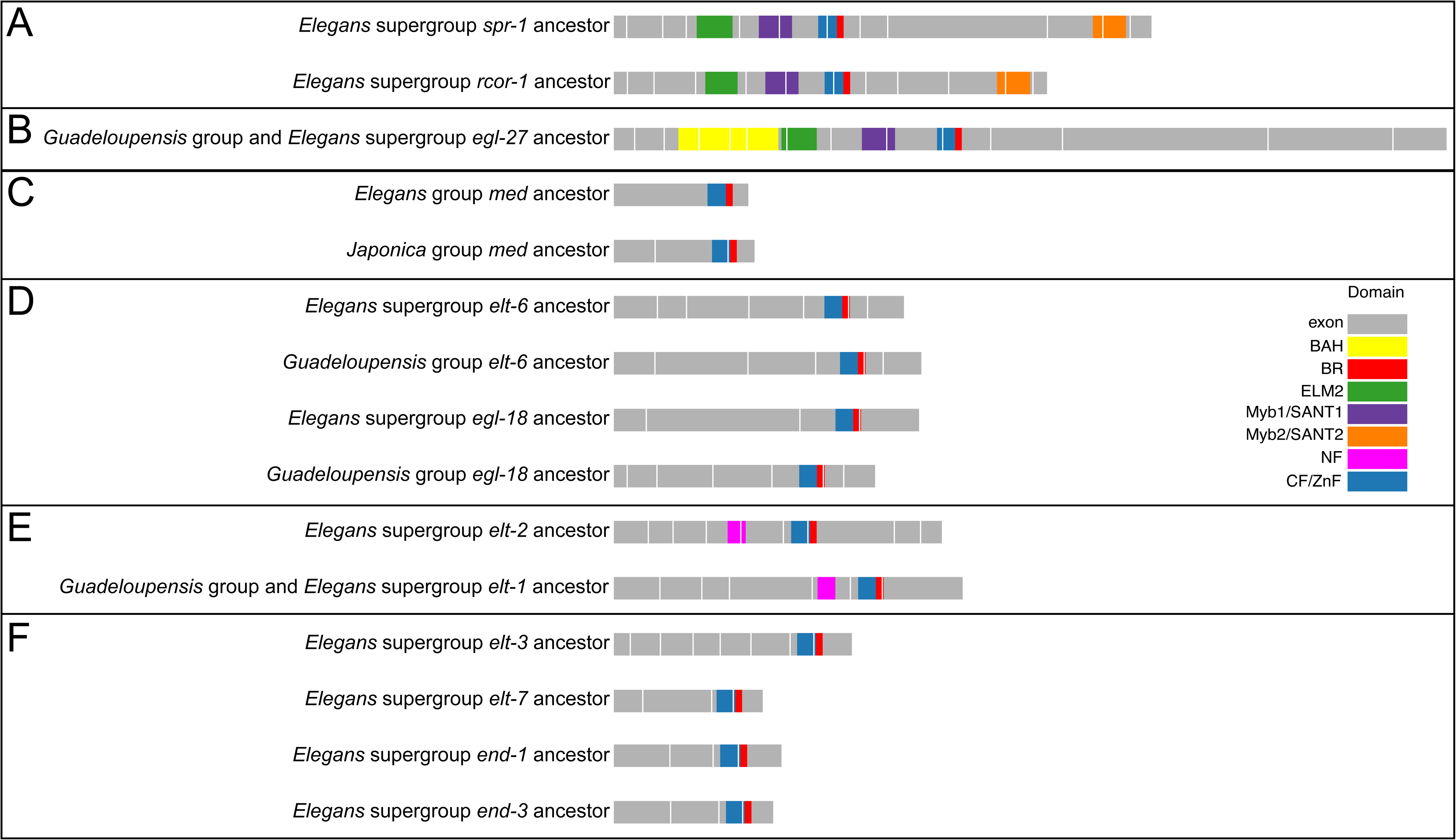
Comparisons of predicted ancestral *Caenorhabditis* GATA-domain-containing gene structures. **(A)** *rcor-1* and *spr-1* predicted *Elegans* supergroup ancestral gene structures. **(B)** Predicted gene structure of the *Guadeloupensis* group and *Elegans* supergroup *egl-27* ancestor. **(C)** Predicted *Elegans* and *Japonica* group ancestral *med* gene structures, respectively. **(D)** *elt-6* and *egl-18* predicted *Elegans* supergroup ancestral gene structures, respectively and *elt-6* and *egl-18* predicted *Guadeloupensis* group ancestral gene structures, respectively. **(E)** Predicted *Elegans* supergroup ancestral *elt-2* gene structure and *Guadeloupensis* group and *Elegans* supergroup *elt-1* ancestral gene structure (which is also representative of the *elt-1 Elegans* supergroup ancestral gene structure). **(F)** *elt-3*, *elt-7*, *end-1*, and *end-3* predicted *Elegans* supergroup ancestral gene structures, respectively. The key to the color coding of the protein domains encoded in the gene structures is shown on the right: exons are shown in gray (with intron positions indicated by white vertical lines); BAH domains are shown in yellow; the BRs of GATA domains are in red; ELM2 domains are shown in green; the Myb/SANT domains nearer the 5’ end of a gene (Myb1/SANT1) are in purple; the Myb/SANT domains nearer the 3’ end of a gene (Myb/SANT2) are in orange; GATA(-like) NFs are in pink; and GATA(-like) CF/ZnFs are in blue.

Due to extensive homology along the entire lengths of the *rcor-1* and *spr-1* sequences (Supp. Fig. 3), we predict that a full-length duplication of *rcor-1* occurred in the *Elegans* supergroup ancestor, thus generating paralogs with the same gene structure as the predicted *rcor-1 Elegans* supergroup ancestor (Fig. 2A). This gene structure consists of 12 exons and has an ELM2 (EGL-27 and MTA1 homology 2) (Solari et al. 1999) domain encoded in exon 4, a Myb/SANT domain encoded in exons 5 and 6 with conservation of the intron location between them, a GATA-like zinc finger motif (ZnF) encoded in exons 6 and 7 with conserved location of the intron between them, and a second Myb/SANT domain encoded in exons 10 and 11 separated by an intron (the location of which is not as highly conserved as the location of the introns associated with the first Myb/SANT domain or the ZnF) (Fig. 2A). ELM2 and Myb/SANT domains do not share homology with GATA factor domains and are known to be involved in transcriptional repression through chromatin regulation (Ding et al. 2003; Boyer et al. 2004; Wang et al. 2006). The presence of one ELM2 domain and two Myb/SANT domains is the signature of genes encoding CoREST proteins (Meier and Brehm 2014). But unlike sequences encoding canonical CoREST proteins, nearly all the *Caenorhabditis* SPR-1 and RCOR-1 orthologs we examined contain a ZnF-like motif (CX_2_CX_16-23_CX_2_C; Supp. Fig. 4) between the two Myb/SANT domains, which, in most cases, has the same length of a typical animal GATA ZnF (i.e., a CX_2_CX_17_CX_2_C motif) (Teakle and Gilmartin 1998; Lowry and Atchley 2000). (These ZnF-like motifs are described in more detail below.) Interestingly, most of the ZnF-like motifs in the SPR-1/RCOR-1 homologs of other nematodes and *Drosophila* have shorter loops of only 14 or 15 residues, suggesting that loop length has changed multiple times as these gene families have evolved. Since the four vertebrate SPR-1/RCOR-1 homologs lack any CX_2,4_CX_14-24_CX_2_C ZnF-like motifs (Wormbase.org), the ZnF-like motif likely arose in invertebrates. In summary, genes coding for the SPR-1 and RCOR-1 proteins in our phylogenetic tree are similar in their organization, including intron splice locations, positions of non-GATA-related domains and of an atypical ZnF (Fig. 2A; Supp. Fig. 2A,B), which supports our hypothesis that these ortholog groups share evolutionary history.

##### *spr-1* and *rcor-1* orthologs contain a not well-conserved GATA-like domain that likely evolved convergently relative to GATA factors

We created a hidden Markov model profile (pHMM) (Eddy 1998) of each of the GATA-like domains in RCOR-1 and SPR-1 proteins (see Materials and Methods; Fig. 3A) and used them to search all the proteins included in our analysis. Other than for SPR-1 orthologs, the SPR-1 pHMM scores were highest for RCOR-1 orthologs and vice versa (Fig. 3B), suggesting a high degree of homology among the GATA domains of these proteins. Moreover, the SPR-1 and RCOR-1 pHMMs had low scores for the other proteins included (Fig. 3B), suggesting lack of conservation with other *Caenorhabditis* GATA domains. This is consistent with the placement of most SPR-1 and all RCOR-1 GATA-like domains closer to each other than to other GATA domains in our GATA domain phylogeny (Supp. Fig. 5).

**Figure 3.**
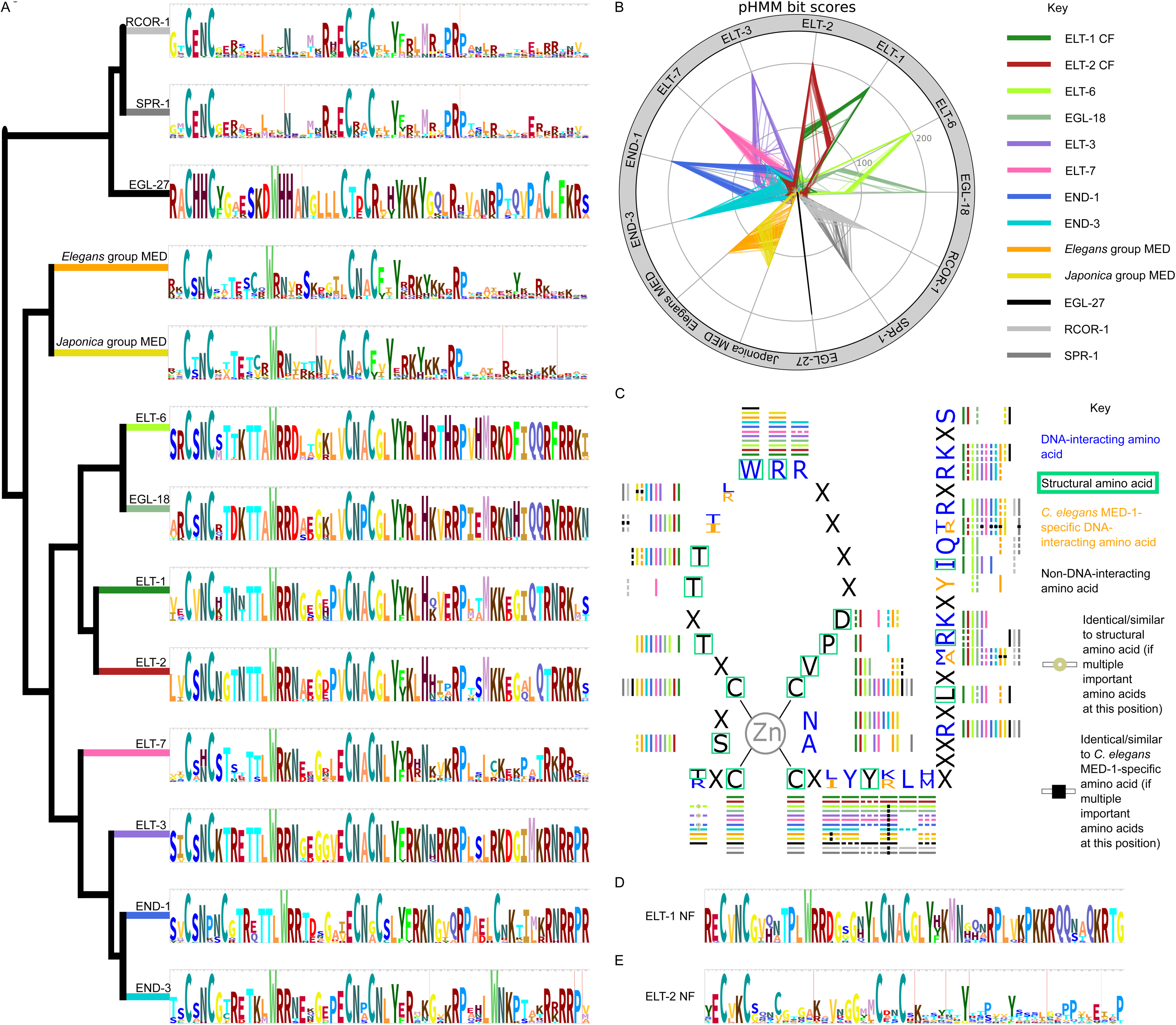
Comparisons of *Caenorhabditis* GATA(-like) DNA-binding domains. **(A)** pHMM amino acid sequence logos, of the CF/ZnF for each ortholog group in the Figure 1A phylogeny (however, the MED ortholog group pHMM is divided into *Elegans* and *Japonica* group MED pHMMs because most of the ZnFs in these two groups have different lengths). The total height of the stack of amino acid(s) at each position represents the total information content at that position (Wheeler et al. 2014). Amino acid(s) with above background frequency scores are shown as subdivisions of the total stack height depending on the probability of that amino acid at that position (Wheeler et al. 2014). The relationships among the ortholog groups in the phylogeny (Fig. 1A) are indicated through the cladogram on the left, the branches of which display the name of the particular ortholog group. **(B)** Radar plot depicting the pHMM bit scores for each of the GATA-domain-containing proteins scored on the 13 pHMMs shown in A. Key to the color-coding by ortholog group is given on the right side of the panel. The scale for the bit scores is depicted by the concentric circles on the figure (see the ELT-6 pHMM radius for numerical values). **(C)** Conservation of important animal GATA factor DNA-binding domain residues in *Caenorhabditis* GATA(-like) DNA-binding domains. Whether or not the residue with the highest probability in an ortholog groups pHMM (Fig. 3A) is the same or is similar (see Supp. Methods) to the indicated residues is denoted by a colored bar to the exterior of the figure. (These bars are color-coded by ortholog group as indicated in the key to the right of B.) Non-dashed colored bars mean that the residues with the highest probability at that position in that ortholog group’s pHMM is identical to the shown residue. If the colored bar is dashed that means that the residues with the highest probability at that position in that ortholog groups pHMM has a similar residue. Chicken GATA-1 (Omichinski et al. 1993), mouse GATA-3 (Bates et al. 2008), human GATA-3 (Chen et al. 2012), and/or *C. elegans* MED-1 (Lowry et al. 2009) residues that interact with DNA are shown in blue. Residues involved in the structural integrity of the chicken GATA-1 bound to DNA (Omichinski et al. 1993) are shown with an aquamarine box around them. *C. elegans* MED-1-specific DNA-interacting residues are shown in orange. Residues not found to interact with DNA in any of these animal GATA factor-DNA structures are shown in black. Some positions have different residues found to be important for the structural integrity and/or DNA-binding in different animal GATA factor-DNA structures (e.g., at the first position a threonine (T) was found to be structurally important in the cGATA-1-DNA structure while an arginine (R) was found to interact with DNA at this position in the mGATA-3-DNA structure). In this case an off-white ring in the middle of the ortholog group colored bar indicates that T has the highest probability of being at this position in this ortholog groups pHMM. Whereas a colored bar without an off-white ring at this position indicates that an R has the highest probability of being at this position in this ortholog groups pHMM. A similar classification, but with a black square, is used for positions where a different residue was found to interact with DNA in the *C. elegans* MED-1-DNA structure compared to at least one of the vertebrate GATA factor-DNA structures. **(D** and **E)** Amino acid sequence logos of the NF pHMMs, of ELT-1 (D) and ELT-2 (E). (Compare these logos based on NFs to those in A based on CF/ZnFs.)

Examination of the SPR-1 and RCOR-1 ZnFs revealed that they all lack an important residue common to GATA factor ZnFs: no proteins in the rcor1 clade have a tryptophan (W) at position 12 in their ZnFs (Fig. 3A; Supp. Fig. 3), which is invariant in most GATA factors (the only exception being some plants, where a methionine (M) occupies this position instead) (Lowry and Atchley 2000). A tryptophan is found at this position in all other *Caenorhabditis* GATA domains included here (Supp. Fig. 7,10,13,14,15). Instead, the residue at position 12 in the ZnFs of *Caenorhabditis* SPR-1 and RCOR-1 orthologs is generally not well conserved. Most of the SPR-1 orthologs we analyzed have glutamic acid (E) at this position, and most RCOR-1 orthologs have an isoleucine (I) (Fig. 3A). A NMR structure of *Gallus gallus* (chicken) GATA-1 (cGATA-1) C-terminal zinc finger (CF) bound to DNA showed that the tryptophan at ZnF position 12 is important for the structural integrity of the ZnF motif (Omichinski et al. 1993). Additionally, a crystal structure of the human GATA-3 (hGATA-3) CF bound to DNA revealed a hydrophobic interaction between the hGATA-3 CF tryptophan at position 12 and a thymine in the DNA (Chen et al. 2012). The function of this tryptophan is likely related to its large size and hydrophobicity. The residues occupying position 12 the ZnFs of SPR-1 and RCOR-1 proteins – glutamic acid and isoleucine – are both smaller residues than tryptophan. Isoleucine is hydrophobic and glutamic acid is negatively charged. A few other canonical animal GATA factor residues are also missing from rcor1 clade GATA domains, most of which are important for the structure or DNA binding of this domain and are found in most *Caenorhabditis* GATA factors. rcor1 clade proteins lack a threonine at the fifth position after the second cysteine, lack arginines at the 9^th^ and 10^th^ positions after the second cysteine, lack a glycine at the 14^th^ position after the second cysteine, and lack an asparagine after the third cysteine (Fig. 3A). These non-canonical residues likely alter the ZnF DNA-binding structure and possibly the binding specificity associated with SPR-1 and RCOR-1 proteins compared to proteins comprising more canonical GATA DNA-binding domains. In fact, there is no evidence that any *Caenorhabditis* SPR-1 or RCOR-1 orthologs bind DNA at all. This domain may instead be used for protein-protein interactions, the structural integrity of the protein, or not at all.

We compared the residue with the highest probability at each position in each pHMM to the residues known to be important for the protein structure and/or DNA binding of animal GATA factors bound to DNA (Omichinski et al. 1993; Bates et al. 2008; Lowry et al. 2009; Chen et al. 2012). These important GATA factor residues are shown in Figure 3C. The RCOR-1 and SPR-1 pHMMs contain the least number of these residues, including one *C. elegans* MED-1-like residue that is not conserved in either of the MED pHMMs (Fig. 3A,C; see below). The RCOR-1 and SPR-1 pHMMs share a similar number of conserved structural and/or binding residues in their GATA-like domains with EGL-27 pHMMs, which is fewer than all other GATA domain pHMMs examined in this study (Fig. 3A,C; see below). Moreover, most of the conserved residues in rcor1 domains are different from those in EGL-27 domains (Fig. 3A,C; see below), supporting their non-adjacent placement of their GATA-like domains in our GATA domain tree (Supp. Fig. 5). If the GATA-like domains in RCOR-1 and SPR-1 shared a common ancestor with canonical GATA factors, their domains must have diverged extensively. Alternatively, the more likely scenario is that these motifs evolved convergently.

##### *spr-1* moved to a new chromosome around the time the ancestral rcor-1 duplicated in the ancestor of the *Elegans* supergroup

We never found *rcor-1* and *spr-1* orthologs on the same piece of assembled genomic DNA (Supp. Fig. 6). In each of the six species in this study for which there was chromosome level genome resolution available (as opposed to only scaffolds and contigs), the *rcor-1* and *spr-1* orthologs were found on different chromosomes (1 and 5, respectively; an additional *spr-1* paralog is present on the X chromosome in *C. inopinata*). To expand our analysis of the chromosome locations of GATA-domain-containing proteins throughout the genus, we assigned scaffolds or contigs to chromosomes based on the *C. elegans* assembly. For each GATA-domain-containing protein on a scaffold or contig in our dataset, we compiled a list of its neighbors, used BLASTp to find their closest homolog in *C. elegans*, and assigned the scaffold or contig to the most common chromosome among these homologs (Fig. 4; see Materials and Methods). Though there have likely been interchromosomal rearrangements during the evolution of this genus, intrachromosomal rearrangements are more frequent (Stein et al. 2003; Teterina et al. 2020) and this is even the case between *C. elegans* and *C. bovis*, a basal species, which mostly have orthologs on the same chromosomes (Stevens 2020).

**Figure 4.**
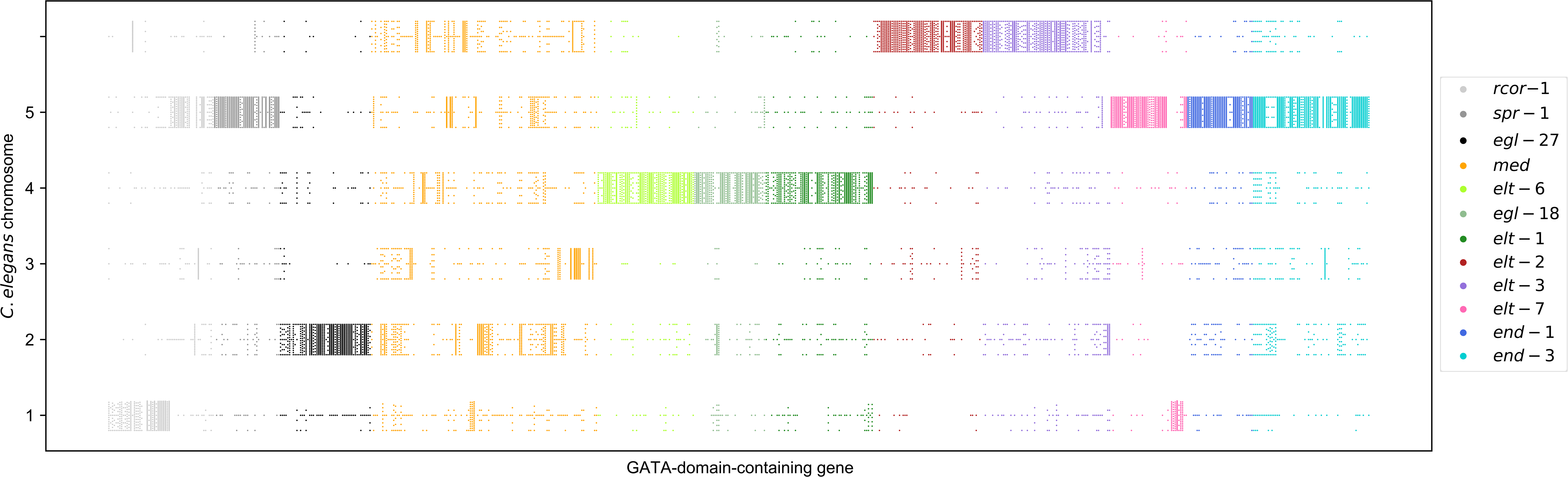
Chromosome assignment for GATA-domain-containing genes on scaffolds or contigs. To expand our analysis of the chromosome locations of GATA-domain-containing proteins throughout the genus, we assigned scaffolds or contigs to chromosomes based on the *C. elegans* assembly. Each dot corresponds to a neighbor (of the gene on the x-axis); the dot location along the y-axis shows the chromosomal location of the *C. elegans* homolog of that neighbor. The color of the dot indicates the ortholog group the gene is assigned to; the key to these color codes is shown on the right. The genes are ordered as in the Figure 1A phylogeny. The numbers and X on the y-axis refer the designations of the six *C. elegans* chromosomes.

In this more detailed dataset we found that while the *Elegans*-supergroup *rcor-1* orthologs were assigned to chromosome 1, non-*Elegans* supergroup *rcor-1* orthologs are assigned to chromosome 5 and so the ancestral gene was probably on chromosome 5 (Fig. 4). The most parsimonious explanation for these results is that during or after the duplication of *rcor-1* and before the split of the *Elegans* supergroup species, a *rcor-1* paralog moved from chromosome 5 to chromosome 1 and this paralog stayed *rcor-1*- like. The other paralog, still on chromosome 5, is the ancestor of extant *spr-1* orthologs. Additionally, this data also suggests that a *C. inopinata*-specific *spr-1* duplication may have occurred interchromosomally or that movement between chromosomes occurred after a tandem duplication.

##### relaxed selection not detected on rcor1 clade paralogs

To test whether the intensity of selection changed after the duplications in the rocr1 clade, we used the RELAX hypotheses testing framework (Wertheim et al. 2015). RELAX compares two sets of branches in a phylogeny and evaluates whether the data is better fit by a single distribution of a few dN/dS rate categories among all branches or by different distributions for each set of branches where the rate categories in one are related to the rate categories in the other by an exponentiation factor (k). We tested whether the *spr-1* orthologs had experienced relaxed selection compared to the *Elegans* supergroup *rcor-1* orthologs and found this to not be the case (p=0.061; k=0.94) (Fig. 5). Additionally, we tested whether *Elegans* supergroup *rcor-1* orthologs had experienced relaxed selection compared to non-*Elegans* supergroup *rcor-1* orthologs. We also did not find significant evidence for this (p=0.17 k=0.94) (Fig. 5).

**Figure 5.**
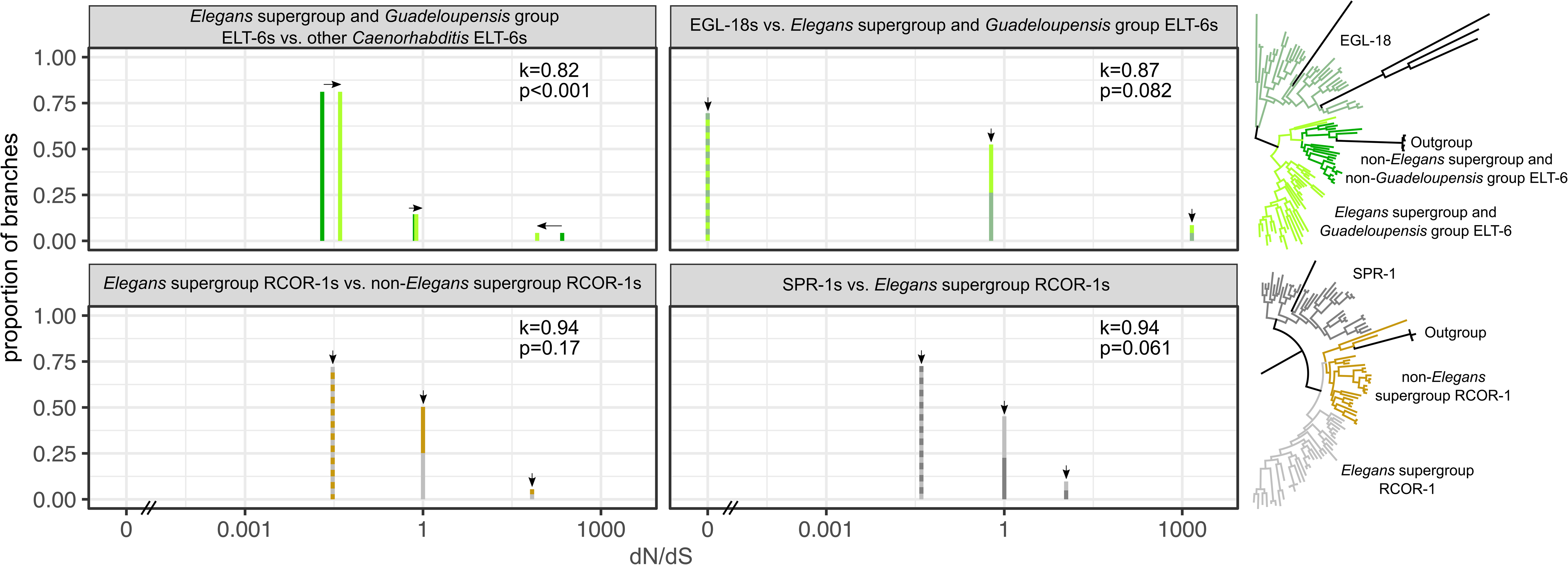
Comparison of selection intensity along RCOR-1 and SPR-1 branches after *rcor-1* duplication and along elt6 clade branches after the divergence of *elt-6* and *egl-18*. The RELAX test was used to test for changes in intensity of selection imposed on the rcor1 and elt6 clade paralogs after their respective ancestral orthologs duplicated. Phylogenetic tree branches used for comparisons are color-coded as per the phylogenetic tree depicted on the right side of the figure (i.e., SPR-1 branches are dark gray, *Elegans* supergroup RCOR-1 branches are in light gray, non-*Elegans* supergroup RCOR-1 branches are in gold, EGL-18 branches are sea green, *Elegans* supergroup/*Guadeloupensis* group ELT-6 branches are in lime green, and other *Caenorhabditis* ELT-6 branches are in dark green). The dN/dS is depicted on the x-axis (the scale of which is the same for all panels). The proportion of branches in each of the three dN/dS rate categories per test is depicted on the y-axis (the scale of which is the same for both panels). The top left panel depicts the RELAX test results comparing selection on *Elegans* supergroup RCOR-1 branches (light gray) to that on SPR-1 branches (dark gray). The bottom left panel depicts the RELAX results comparing selection on *Elegans* supergroup RCOR-1 branches (light gray) to that on non-*Elegans* supergroup RCOR-1 branches (gold). The top right panel depicts the RELAX test results comparing selection on *Elegans* supergroup ELT-6 branches (lime green) to that on EGL-18 branches (dark sea green). The bottom right panel depicts the RELAX results comparing selection on *Elegans* supergroup ELT-6 branches (lime green) to that on non-*Elegans* supergroup ELT-6 branches (dark green). The exponentiation factors (k) and p-values for differences in dN/dS rate category distributions for each test are shown in the top right corner of each panel. Arrows in the panels indicate the direction of selection pressure; arrows pointing towards a dN/dS ratio of one indicate relaxed selection, those pointing toward values less than one indicate increasing negative selection, and those pointing to values greater than one indicate increasing positive selection.

#### EGL-27 ortholog group

##### EGL-27 orthologs form a monophyletic clade that is more likely to share a more recent common ancestor with orthologs of the rcor1 clade than with canonical GATA factors

*C. elegans* EGL-27 functions in embryonic patterning by controlling cell polarity, migration, and fusion (Ch’ng and Kenyon 1999; Herman et al. 1999; Solari et al. 1999). We identified at least one EGL-27 ortholog in 59 of 60 species included in our phylogenetic analysis (Fig. 1; Supp. Table 2), suggesting that *egl-27* originated before split of *Caenorhabditis* and *Diploscapter* and may have been lost in *C. virilis*. EGL-27 orthologs form a well-supported (100% ultrafast bootstrap support (Minh et al. 2013; Hoang et al. 2018)) group in our phylogeny (Fig. 1A). These orthologs form a tight, distinct cluster closer to the rcor1 clade than to any other clade or ortholog group (Fig. 1A). Like the rcor1 clade genes, they contain ELM2 and Myb/SANT domains (Fig. 2B) as well as an atypical GATA ZnF (Fig. 3A-C). And like the rcor1 clade orthologs, EGL-27 orthologs share homology with non-GATA factor proteins in other organisms (Wormbase.org), such as the human arginine-glutamic acid dipeptide repeats protein encoded by the RERE gene (Wang and Tsai 2008), which has the same domains and similar organization as *Caenorhabditis* EGL-27s but with an additional Atrophin-1 domain (Wang and Tsai 2008). Atrophin proteins repress transcription through recruitment of histone modifiers (Wang et al. 2006; Wang et al. 2008) but can be transcriptional activators in other contexts (Shen et al. 2007; Vilhais-Neto et al. 2010). Taken together, these data suggest that the EGL-27 orthologs likely share a more recent common ancestor with genes of the rcor1 clade than with any of the other genes we examined in this study.

##### *egl-27* genes are longer and are divergent from those in other ortholog groups

We compared extant *egl-27* gene structures and made predictions about their ancestral gene(s) (Supp. Fig. 2C). For example, based on the large numbers of exons in the extant *egl-27* genes (more than in any other gene family included in this study) we predict that the ancestral *Guadeloupensis* group and *Elegans* supergroup *egl-27* had 14 exons (Fig. 2B). Moreover, EGL-27 protein sequences are substantially longer than all other protein sequences in our study (Fig. 2; Supp. Fig. 2). Only the EGL-27 orthologs contain a bromo-adjacent homology (BAH) (Nicolas and Goodwin 1996) domain (Fig. 2; Supp. Fig. 2). BAH domains, which are often associated with gene silencing through protein-protein interactions (Callebaut et al. 1999), are large. In the case of EGL-27 orthologs the domain usually spans 4 exons (Fig. 2B; Supp. Fig. 2C). Additionally, EGL-27 orthologs only have a single Myb/SANT domain whereas most rcor1 clade orthologs have two (Fig. 2A,B; Supp. Fig. 2A-C). *egl-27* orthologs are also distinguished by a conserved splice site immediately following the first 21 nucleotides coding for their ELM2 domain, a somewhat conserved splice site immediately following the first 24 or 102 nucleotides coding for their Myb/SANT domain, and a conserved splice site immediately following the first 22 nucleotides coding for their atypical GATA ZnFs (see below) (Fig. 2B; Supp. Fig. 2C). These conserved splice sites are not found in any of the other GATA-domain-containing proteins included in this study (Supp. Fig. 2), which further supports the separate evolutionary history of *egl-27* orthologs. Overall, the EGL-27 orthologs are encoded by large genes, with conserved splice sites at different locations from those conserved in the other ortholog groups in this study, and have three non-GATA factor domains, two of which are also found in rcor1 clade genes. These results support their placement close to, but not within, the rcor1 clade in our phylogeny (Fig. 1A).

##### EGL-27 orthologs contain atypical GATA DBDs that are likely on a divergent evolution trajectory in relation to those of other Caenorhabditis GATA-domain-containing proteins

We created a pHMM (Eddy 1998) of the atypical GATA domain in EGL-27 orthologs (Fig. 3A) and used it to search against all the proteins in our analysis (see Materials and Methods). The EGL-27 profile only scored EGL-27 orthologs well (Fig. 3B), which indicates the uniqueness of this GATA-like domain in *Caenorhabditis*. There are two primary reasons EGL-27 ZnFs are classified as GATA factors. One is due to the conserved tryptophan (W) at position 12 in their ZnFs and the other is their similarly spaced cysteines, i.e., CX_2_CX_16_CX_2_C (Fig. 3A; Supp. Fig. 7; Supp. Fig. 4; prosite.expasy.org) (Lowry and Atchley 2000). All animal GATA factors have a tryptophan at position 12; however, most of their cysteine pairs are 17 residues apart, as opposed to the 16 in EGL-27 orthologs (Teakle and Gilmartin 1998; Lowry and Atchley 2000; Gillis et al. 2008; Maduro 2020). This shorter ZnF loop in *Caenorhabditis* EGL-27 orthologs is why their ZnFs are classified as atypical (prosite.expasy.org) (Lowry and Atchley 2000). Only in three *Japonica* group sister species, *C. nouraguensis*, *C. becei*, and *C. yunquensis*, do EGL-27s have the typical 17 residues in their ZnF loops (Supp. Fig. 7), which suggests that a 3 base pair (bp) insertion into the *egl-27* ZnF coding sequence or an alternative splice site likely occurred in their most recent common ancestor. Interestingly, *Diploscapter* EGL-27 paralogs, EGL-27 orthologs from two Clade II nematodes, EGL-27 orthologs from a Clade III nematode, and the mouse and human RERE proteins all have ZnF loops that are 18 residues long, whereas the EGL-27 ortholog from the Clade V *Pristionchus pacificus* nematode has a ZnF loop with 19 residues. *Drosophila* species appear to lack an EGL-27 ortholog altogether (Wormbase.org) (Haag et al. 2018). Overall, the loop size of EGL-27/RERE orthologs has likely changed multiple times in different lineages, and the orthologous gene may have been lost from the ancestor of the *Drosophila* lineage. Although there are a few exceptions (see details in the elt1/2 clade and MED ortholog group sections below), since most other *Caenorhabditis* GATA factor orthologs have ZnFs with the typical animal loop length of 17 residues (Supp. Fig. 4), the atypical length of most EGL-27 ortholog ZnF loops is more consistent with a separate evolutionary history thus supporting this clade’s placement in our phylogeny more distant from the clades of more canonical GATA factors (Fig. 1A).

There are 21 residues in EGL-27 ZnFs and BRs that distinguish them from typical *Caenorhabditis* GATA domains. Ten of them are also found in the human and mouse RERE proteins, which have identical GATA DBDs (Supp. Fig. 8), suggesting that these 10 have been highly conserved since the vertebrates and invertebrate ancestors diverged. The 21 EGL-27-specific ZnF and BR residues are: histidines (H) at positions three and four (note that position numbering in Supplemental Figure 8 starts at zero), a tyrosine (Y) or a phenylalanine (F) at position six, aspartic acid (D) at position 12, histidines at positions 14 and 15, a threonine (T) at position 25, an aspartic acid or a glutamic acid (E) at position 26, an arginine (R) at position 28, usually a leucine (L) at position 37, usually a non-proline (non-P) at position 39, usually an alanine (A) or a valine (V) at position 41, usually an asparagine (N) at position 42, a proline at position 44, usually a threonine at position 45, usually a glutamine (Q) at position 46, a proline at position 48, usually an alanine at position 49, a cysteine (C) at position 50, a leucine at position 51, and a phenylalanine at position 52 (Supp. Fig. 8). Similar residues (Henikoff and Henikoff 1992) are rarely found in these positions in canonical *Caenorhabditis* GATA domains (Fig. 3A,D,E; Supp. Fig. 10,13,14,15). The few GATA factors that do have some of these residues do not have more than three of them (Supp. Fig. 10,13,14,15). These minor similarities between canonical GATA ZnFs and EGL-27 ZnFs are likely due to convergent evolution. 11 of these 21 EGL-27-specific ZnF residues are known to be important for the structure and/or binding function of the cGATA-1 DBD or the DNA binding of mouse and human GATA-3s (Omichinski et al. 1993; Bates et al. 2008; Chen et al. 2012) and so may affect the DNA-binding kinetics or specificity of EGL-27 orthologs. For example, the *C. elegans* EGL-27 ortholog binds not only to a GATA site (GATAAG) but also to the non-GATA sites GAGAAG and RGRMGRWG (Xu and Kim 2012).

The number of binding or structural residues that are identical or similar between canonical animal GATA domains and those in the EGL-27 and rcor1 clade orthologs are approximately the same (Fig. 3C; see above), and there are fewer similarities compared to the more typical GATA factor ortholog groups in this study (Fig. 3C). Additionally, the similarities in EGL-27 orthologs are different from the similarities in rcor1 clade orthologs (Fig. 3C), suggesting that their GATA-like domains likely do not share a recent common ancestor. The shared ELM2 and Myb/SANT domains rather than the GATA domain may be the shared features between EGL-27 and rcor1 clade orthologs that place them on a closer branch to each other than to GATA factors in our phylogeny (Fig. 1A), especially since these GATA domains do not group adjacently in our GATA domain phylogeny (Supp. Fig. 5). The EGL-27 GATA domains are highly conserved across the genus (Fig. 3A; Supp. Fig. 5), suggesting they have experienced strong negative selection. Moreover, many of the non-canonical features of the EGL-27 GATA-like motifs are conserved among vertebrate RERE proteins (Supp. Fig. 8) suggesting that EGL-27 ZnF and BRs evolved separately from canonical GATA factor DBDs at least since the split between vertebrate and invertebrates and perhaps are strictly convergent.

##### Most *egl-27* orthologs are not syntenic with any other GATA-domain-containing proteins

We only found two *Caenorhabditis* species with their *egl-27* ortholog syntenic with any GATA factors or rcor1 clade orthologs included in this study (Supp. Fig. 9). The first is in *C*. sp. *44*, where we found that the *egl-27* ortholog is 12 Mb from the *rcor-1* ortholog (Supp. Fig. 9). The second is in *C. tropicalis* where the *egl-27* ortholog is on the same scaffold as the *elt-2*, *elt-3*, and *egl-18* orthologs, a truncated *elt-6* ortholog, and three truncated *med* orthologs (Supp. Fig. 9). This is unexpected, since most of these other orthologs are located on three different chromosomes in other species (Fig. 4). In the few species with available chromosome-level assemblies (which does not include *C. tropicalis*) *egl-27* orthologs are found on chromosome 2, *elt-2* and *elt-3* orthologs are located on the X chromosome, and *egl-18* and *elt-6* orthologs are found on chromosome 4. Interestingly, we found that the close neighbors of *C. tropicalis egl-*27, *egl-18*, *elt-2*, and *elt-3* orthologs in *C. elegans* are mainly on chromosome 1, 4, X, and X, respectively, which are the same chromosomes these orthologs are found on in species with chromosome-level assemblies (Fig. 4). This suggests that interchromosomal rearrangements of at least kilobase long multi-gene sequences from at least three chromosomes may have occurred in *C. tropicalis*. It will be interesting to re-examine synteny in *C. tropicalis* once its genome has been more fully assembled. Until then, our data suggest that *C. tropicalis* has undergone substantial chromosome rearrangements as compared to most of the other species included in this study.

### Canonical *Caenorhabditis* GATA factors

#### elt6 clade

##### *egl-18* originated through a duplication of elt-6 that occurred in the common ancestor of the *Elegans* supergroup and *Guadeloupensis* group species

Our phylogeny places the ELT-6 and EGL-18 (also known as ELT-5) ortholog groups in a well-supported (100% ultrafast bootstrap support (Minh et al. 2013; Hoang et al. 2018)) monophyletic clade (Fig. 1A). A previous phylogenetic study of *C. elegans* and *C. briggsae* also placed ELT-6 and EGL-18 orthologs in a well-supported monophyletic clade (Gillis et al. 2008). In *C. elegans*, ELT-6 and EGL-18 function redundantly in seam cell specification (Koh and Rothman 2001), larval seam cell maintenance (Gorrepati et al. 2013), male tail morphogenesis (Nelson et al. 2011), and inhibition of cell fusion during vulval development (Koh et al. 2002). However, ELT-6 and EGL-18 do not necessarily contribute equally to these phenotypes (Nelson et al. 2011; Gorrepati et al. 2013). While the functions of ELT-6 and EGL-18 orthologs in other *Caenorhabditis* species have not been studied, conservation in protein sequence among the ELT-6 and EGL-18 orthologs in this study (Supp. Fig. 10) make it likely that their functions are also conserved. Because EGL-18 orthologs are only found in *Elegans* supergroup and *Guadeloupensis* species, whereas ELT-6 orthologs are present throughout the *Caenorhabditis* genus as well as in the *Diploscapter* species included in our study (Fig. 1), we conclude that an *elt-6* duplication produced *egl-18* in the ancestor of the *Elegans* supergroup and *Guadeloupensis* group.

##### The *Elegans* supergroup ancestral egl-18 likely lost three introns while the *Guadeloupensis* group ancestral elt-6 likely lost a single exon

Most of the singleton *elt-6* orthologs (30 of 40) have seven exons and code for a single GATA ZnF within their third to last exon (Supp. Fig. 2D). This gene structure appears to have been conserved among *egl-18* genes in *Guadeloupensis* group species as well, at least in the two species for which genomic sequences are available (Supp. Fig. 2E). This suggests that a full-length *elt-6* duplication produced the *egl-18* ancestor in the *Guadeloupensis* and *Elegans* supergroup ancestor. Interestingly, all 30 singleton *egl-18* orthologs in the *Elegans* supergroup have only four exons, and they also code for a single GATA ZnF in their second to last exon (Supp. Fig. 2E). Yet despite these differences in their gene structures, the protein sequences of ELT-6 and EGL-18 orthologs are of similar lengths (medians of 389 and 420 residues, respectively) and align along their entire lengths (Supp. Fig. 10). These data suggest that the *Elegans* supergroup ancestral *egl-18* lost three introns (Fig. 2D). The locations of the ZnF-coding exons indicate that the *Elegans* supergroup *egl-18* gene lost intron 6, which was the final intron in its *elt-6* ancestor (Fig. 2D). Additionally, *Elegans* supergroup *egl-18s* have large second exons (median 636 nts), which are larger than the combined lengths of exons 2, 3, and 4 of most *elt-6* genes (median total 595.5 nts) (Supp. Fig. 10). This suggests that the *Elegans* supergroup *egl-18* ancestor also lost introns 2 and 3, relative to its *elt-6* ancestor (Fig. 2D). *Guadeloupensis* group *elt-6s* likely also lost intron 2 as compared to most other *elt-6* orthologs in this study (Fig. 2D). *Guadeloupensis* group *elt-6* orthologs also have larger exon 2s (median 383.5 nts) which approximate the combined lengths of exons 2 and 3 (median 371.5 nts) in most *elt-6* genes (Fig. 2D). Our analyses of phylogeny (Fig. 1) and synteny (see below) suggest that the loss of intron 2 in *Elegans* supergroup *egl-18* genes and *Guadeloupensis* group *elt-6* genes occurred convergently, although another possibility is that the *Elegans* supergroup ancestral *egl-18* is more closely related to the *Guadeloupensis* group ancestral *elt-6* than to the *Guadeloupensis* ancestral *egl-18*.

##### GATA domains of elt6 clade orthologs contain a conserved intron in their basic regions

The *elt-6* and *egl-18* orthologs have a conserved intron 24 nucleotides into the basic region (BR), which is directly C-terminal to the ZnF motif (Supp. Fig. 2D,E). This intron location is conserved for all singleton *elt-6* and *egl-18* genes in this study as well as in all singleton *elt-1* CFs apart from the *C. sinica elt-1* CF (Supp. Fig. 2D-F). Eurmsirilerd & Maduro (2020) found this conserved intron location in many, mostly non-*Caenorhabditis*, nematode *egl-18*/*elt-6*, *elt-1*, and Clade I *elt-2* CFs and we extend this finding to *elt-6*, *egl-18*, and *elt-1* orthologs in 53 more *Caenorhabditis* species. This splice site location is also found in vertebrate and some arthropod GATA factors and was likely the splicing location in the bilaterian ancestral GATA factor CF (Gillis et al. 2008). This conserved intron location supports our phylogenetic placement of the ELT-6 and EGL-18 ortholog groups into a monophyletic clade. It also hints at a relationship between the elt6 clade genes and the CFs of genes in the elt1/2 clade.

##### EGL-18 and ELT-6 orthologs have similar GATA DNA-binding domains

We created pHMMs (Eddy 1998) of the ELT-6 and EGL-18 DBDs (Fig. 3A) and used them to search against all the proteins included in our analysis (Fig. 3B; see Materials and Methods). We found the ELT-6 ortholog group pHMM scores higher against EGL-18 orthologs than any others except ELT-6 orthologs and vice versa (Fig. 3B), indicating that the DBDs of these ortholog groups are most like each other. The ELT-6 and EGL-18 orthologs uniquely and exclusively have an alanine (A) at position seven in their ZnF loops (Fig. 3A). This alanine is also conserved in all the ZnFs of ELT-6/EGL-18 orthologs identified across the Nematode phylum (Eurmsirilerd and Maduro 2020). An NMR structure of cGATA-1 DBD bound to DNA showed that the leucine (L) at position seven in the cGATA-1 ZnF loop interacted with DNA in the major groove (Omichinski et al. 1993), and most other *Caenorhabditis* GATA factors have a leucine conserved at this position as well (Fig. 3). The *Aspergillus nidulans* AreA GATA factor also has a leucine at this position, as do most animal and some other fungi GATA factors (Teakle and Gilmartin 1998). Mutations of this leucine to a valine (V) significantly alter the binding specificity and affinity of AreA *in vivo* (Ravagnani et al. 1997) and *in vitro*, such that it binds better to TGATA better than to CGATA DNA sites (Starich et al. 1998). A mutation in AreA at this same site to a Methionine (M) has weaker but opposite effect (Arst and Scazzocchio 1975). Because both valine and alanine are smaller than leucine, whereas methionine is larger, the alanine residue found in ELT-6 and EGL-18 orthologs may also increase specificity for TGATA over CGATA DNA sites and likely alters the ZnF interactions with DNA in some way that could change the DNA-binding kinetics of these orthologs.

We also compared the residues of highest probability at each position in both the ELT-6 and EGL-18 pHMMs to the residues found to be important for structure or DNA binding in animal GATA factors bound to DNA (Fig. 3C) (Omichinski et al. 1993; Bates et al. 2008; Lowry et al. 2009; Chen et al. 2012). The ELT-6 pHMM has 19 residues identical to and one residue similar to the 24 residues found to interact with DNA. Additionally, the ELT-6 pHMM has 13 residues identical to and two residues similar to the 18 residues found to be important for the structural integrity of the cGATA-1 DBD (Fig. 3C). The EGL-18 pHMM is very similar to the ELT-6 pHMM with 18 residues identical to and two residues similar to the 24 residues important for DNA binding and 13 residues identical to and one residue similar to the 18 residues important for the DBD structure (Fig. 3C). Of the twelve ortholog groups in our phylogeny, the ELT-6 and EGL-18 pHMMs contain the second and third most conserved functional residues, respectively, suggesting that their DBDs are likely under similar levels of stabilizing selection.

##### *egl-18* likely originated from a tandem duplication of elt-6, and dicistron transcription of these genes may be conserved throughout the *Elegans* supergroup and *Guadeloupensis* group species

*elt-6* and *egl-18* are adjacent to each other on the same chromosome/scaffold/contig in 30 of the 31 species where we could confidently identify orthologs of both genes (Supp. Fig. 11). Moreover, in all these species *egl-18* orthologs are found upstream of *elt-6* orthologs and on the same strand and thus transcribed in the same direction (Supp. Fig. 11). In *C. elegans, egl-18* and *elt-6* are sometimes transcribed together as a dicistron (Koh and Rothman 2001), and since this operon-like structure is conserved in many of the species in this study, these additional species may also express these genes via a dicistron. The tight synteny between *elt-6* and *egl-18* orthologs supports our hypothesis that a duplication of *elt-6* produced the *egl-18* ancestor, and this ancient duplication was probably a tandem duplication.

##### Selection was relaxed on elt-6 paralogs relative to ancestral singleton elt-6

We tested if *elt-6* orthologs experienced different degrees of selection pressures after gene duplication in the *Elegans* supergroup/*Guadeloupensis* group ancestor. We found significant relaxed selection on *Elegans* supergroup/*Guadeloupensis* group *elt-6* orthologs compared to that of the other *Caenorhabditis elt-6* orthologs (p<0.001; k=0.82) (Fig. 5). We did not find significant evidence for selection differences between *Elegans* supergroup/*Guadeloupensis* group *elt-6*s compared to that on their *egl-18* paralogs (p=0.082; k=0.87) (Fig. 5).

#### ELT-2 ortholog group

The ELT-2 ortholog group includes both the *C. elegans* ELT-2 and ELT-4 proteins; ELT-4 is on a branch near *C. elegans* ELT-2, but it does not group adjacent to it (Fig. 1). *C. elegans* ELT-4 has no known function and is thought to be a pseudogene that resulted from a *C. elegans*-specific *elt-2* duplication (Fukushige et al. 2003). We did not identify any *elt-4* orthologs or even other *elt-2* duplications in any of the 57 other *Caenorhabditis* species, supporting the hypothesis that a duplication of *elt-2* within *C. elegans* produced *elt-4* and that its position away from *C. elegans elt-2* in our tree reflects a loss of selective constraint on its sequence.

#### elt1/2 clade

##### ELT-1 and ELT-2 orthologs group adjacently forming an elt1/2 clade with ancient origins

The elt1/2 clade consists of the ELT-1 and ELT-2 ortholog groups, which cluster adjacently forming a well-supported monophyletic clade (100% ultrafast bootstrap support (Minh et al. 2013; Hoang et al. 2018)) (Fig. 1A). We found at least one *elt-1* ortholog and one *elt-2* ortholog in every *Caenorhabditis* and *Diploscapter* species included in this study (Fig. 1), and *elt-1* and *elt-2* orthologs have also been found in all extant nematodes with fully sequenced genomes (Eurmsirilerd and Maduro 2020). The ancestors of these genes were probably present at the beginning of the nematode phylum. In *C. elegans,* ELT-1 and ELT-2 orthologs function in different germ layers: ELT-1 is involved in the specification and differentiation of hypoderm precursors (Page et al. 1997; Gilleard and McGhee 2001), while ELT-2 is involved in differentiation and maintenance of endoderm cells (Fukushige et al. 1998). Additionally, the gene structures of *elt-1* and *elt-2* orthologs are different (Supp. Fig. 2F,G; see below) (Eurmsirilerd and Maduro 2020) and we found no cases in which *elt-2* and *elt-1* orthologs were syntenic (Supp. Fig. 12).

ELT-1 and ELT-2 are unusual in *Caenorhabditis* because they both have two zinc fingers (Fig. 2E; Supp. Fig. 2F,G; Supp. Fig. 13). Their CFs are similar (Fig. 3A; Supp. Fig. 13; see below), but their N-terminal ZnF (NF) sequences have diverged (Fig. 3D,E; Supp. Fig. 13; see below). In fact, it appears that the ELT-2 NF has experienced relaxed selection to the point where most ELT-2 NFs are barely recognizable other than their two cysteine pairs (Fig. 3E; Supp. Fig. 13; see below). Vertebrate GATA factors have two canonical GATA ZnF motifs whereas invertebrate, plant, and fungi GATAs have either one or two motifs (Reyes et al. 2004). We therefore hypothesize that the shared ancestry of the elt1/2 clade goes back further than for any of the other well-supported clades in our phylogeny and that the sequences and likely the function(s) of the NFs of these GATA factors have diverged extensively since their most recent common ancestor.

##### *elt-1* and *elt-2* ortholog GATA domains are encoded differently

Even though many *elt-1* and *elt-2* orthologs have nine exons and code for proteins of similar length, there are conserved gene structure features that distinguish the genes of these two ortholog groups from each other (Supp. Fig. 2F,G). Our predicted ancestral elt1/2 clade gene structures provide a visual summary of the differences (Fig. 2E; Supp. Fig. 2F,G). Most of the ZnF motifs are coded in different exons and have different splice sites (Supp. Fig. 2F,G). For example, the NF in the *Elegans* supergroup ancestral *elt-2* was coded in exon 4 whereas it was coded in exon 5 in the ancestral *elt-1* of the *Elegans* supergroup and even one split earlier in the ancestor of the *Guadeloupensis* and the *Elegans* supergroup (Fig. 2E). Moreover, the introns in extant *elt-1* orthologs are nearly always longer than the introns in *elt-2* orthologs, and the gene structures of the latter are relatively compact (data not shown). *elt-2* orthologs all have a conserved intron located just upstream of the last seven nucleotides comprising their CFs (Supp. Fig. 2G; Fig. 2E). This feature is shared with the elt3 clade (see below) and *Japonica* group *meds* (see above), and is found in most nematode *elt-2* and *elt-3* orthologs (Eurmsirilerd and Maduro 2020) and most previously examined *Caenorhabditis end-1*, *end-3*, *elt-2*, and *Japonica* group *med* orthologs (Maduro 2020). *elt-2* NFs have a less well conserved intron position located upstream of the last 19 nucleotides comprising that motif that is not found in any of the other genes in this study (Supp. Fig. 2G; Fig. 2E). *elt-1* orthologs have a conserved intron position located just after the first 24 nucleotides comprising their CF basic regions, which is also found in elt6 clade genes (see above), and their NFs have a conserved intron position located 60 nucleotides after the end of the ZnF sequence (Supp. Fig. 2F; Fig. 2E). These two splice site locations are likely conserved from the bilaterian ancestral GATA factor (Gillis et al. 2008) and were previously found in some nematode *elt-1* and *elt-6* orthologs (Eurmsirilerd and Maduro 2020). This NF intron location is not found in any of the other genes in this study (Supp. Fig. 2). Additionally, the spacing between the NF and CF in *elt-1* orthologs is highly conserved. All singleton *elt-1* orthologs outside the *Elegans* supergroup have 29 residues between their NF and CF motifs, while in most singleton *Elegans* supergroup *elt-1* genes that spacing is 30 residues (Supp. Fig. 2F). Twenty-nine residues between the N- and CFs is also the predicted state of the bilaterian ancestral GATA factor (Gillis et al. 2008), suggesting that this spacing represents the ancestral state and that the ancestor *elt-1* of the *Elegans* supergroup acquired another residue between its ZnFs. On the other hand, the numbers of residues between the NFs and CFs of singleton *elt-2* orthologs is more variable, ranging from 36 to 89 residues (Supp. Fig. 2G). Overall, the divergent organization of these ZnF domains in *elt-1* and *elt-2* orthologs indicate divergent evolutionary paths since they last shared a common ancestor.

##### ELT-1 and ELT-2 orthologs have C-terminal GATA domains that are similar to each other and to the GATA factors of arthropod and vertebrate species

The CF GATA domains of ELT-1 and ELT-2 orthologs share long stretches of common residues (Fig. 3A; Supp. Fig. 13), and this pattern holds across the nematodes (Eurmsirilerd and Maduro 2020). Furthermore, the sequences of their CFs are more similar to those of arthropod and vertebrate GATA CFs than to the single ZnF domains of other *Caenorhabditis* GATA-motif-containing proteins (Fig. 3C; data not shown). The similarities between ELT-1 and ELT-2 CFs are captured by our pHMMs in which each of these proteins’ pHMMs scores the other ortholog’s CF sequences second highest after that of their own ortholog’s CFs (Fig. 3B). Moreover, the ELT-1 and ELT-2 CFs group adjacently in our GATA domain tree (Supp. Fig. 5). The highly conserved PVCNACGLY[FY]KLH sequence, located at positions 20-25 of the CF and followed by the first seven residues of the BR (Fig. 3A), illustrates the similarities between the ELT-1 and ELT-2 CF domains. Structures of vertebrate GATA factors showed that this sequence encodes for the second anti-parallel beta-sheet and all but the last residue of the alpha helix of some canonical GATA factors (Omichinski et al. 1993; Bates et al. 2008; Chen et al. 2012). This sequence is found in all singleton ELT-1 and ELT-2 orthologs in this study, with the single exception of the *C. monodelphis* ELT-2, which has a threonine (T) instead of the proline (P) at the start (Supp. Fig. 13). This particular sequence is also highly conserved in the CFs of most arthropod and vertebrate GATA1/2/3 factors and, with only one residue different, in the CFs of vertebrate GATA4/5/6 factors, which have a methionine (M) instead of a phenylalanine (F) or tyrosine (Y) (Teakle and Gilmartin 1998). The CFs of most (107 of 109) singleton ELT-1 and ELT-2 orthologs contain another highly conserved sequence, TTLWRRN, in positions 11-17 (Fig. 3A), which is also highly conserved in ELT-3 orthologs and in arthropod and vertebrate GATA factors CFs (Teakle and Gilmartin 1998). The TTLWRRN sequence is also present in two non-sister species END-1s, but this is more likely due to convergent evolution since most END-1s have the sequence TTLWRRT at this location (Fig. 3A; Supp. Fig. 13). No other *Caenorhabditis* GATA-domain-containing proteins have TTLWRRN in their ZnF sequences (Supp. Fig. 4; Supp. Table 1). The mostly invariant sequence conservation between the CF domains of ELT-1 and ELT-2 orthologs is evidence of their shared ancestry and suggests strong functional constraint of these domains through negative selection.

We also compared the more probable residue at each position in both the ELT-1 and ELT-2 CF pHMMs to the residues known to be important for the structure or DNA binding in animal GATA factors (Fig. 3C) (Omichinski et al. 1993; Bates et al. 2008; Lowry et al. 2009; Chen et al. 2012). The ELT-1 CF pHMM contains 22 of the 24 residues important for DNA interactions plus one similar residue (Fig. 3C). Additionally, this pHMM contains 13 of the 18 structurally important residues plus two similar residues (Fig. 3C; see Materials and Methods). The ELT-2 pHMM has 16 identical and three similar residues to the 24 residues important for DNA binding and 11 identical and two similar residues to the 18 structurally important residues (Fig. 3C). Of the twelve ortholog groups in our phylogeny, the ELT-1 CF pHMM contains the most of these highly conserved structural and/or DNA-binding residues, while the ELT-2 CF pHMM has the fourth most. This suggests that ELT-1 orthologs have likely experienced stronger negative selection pressures since the divergence of ELT-2 and ELT-1 paralogs.

##### The N-terminal zinc fingers of ELT-1 and ELT-2 orthologs have diverged

The NFs of *Caenorhabditis* ELT-2 orthologs branch off from a very long internal branch within the ELT-1 NF clade in our GATA domain phylogeny (Supp. Fig. 5). This topology is similar to their nematode-wide pattern (Eurmsirilerd and Maduro 2020), suggesting that the NFs of these two ortholog groups are on different evolutionary trajectories. In fact, the ELT-2 NF is highly diverged from the canonical animal GATA factor NF sequence, containing both fewer conserved residues and a variable ZnF loop length (Fig. 3E; Supp. Fig. 13). The ELT-1 NF pHMM contains 13 of the 24 residues important for CF DNA interactions, 11 of 11 residues found involved in NF DNA binding (two of which are not included in the 13 CF residues), and eight of the 18 structurally important CF residues (Fig. 3C,D). The ELT-2 pHMM contains only one of the 24 residues important for CF DNA binding, zero of 11 residues found involved in NF DNA binding, and only four of the 18 structurally important residues (Fig. 3C,E). Because it cannot independently bind DNA and lacks many conserved residues, the NF of the *C. elegans* ELT-2 ortholog has been suggested to be non-functional and degenerate (Hawkins and McGhee 1995; Eurmsirilerd and Maduro 2020). However, some canonical and highly conserved vertebrate NFs do not bind to DNA independently (Martin and Orkin 1990; Yang and Evans 1992)(Martin and Orkin 1990), and some NFs are known to participate in protein-protein interactions (Tsang et al. 1997; Ono et al. 1998; Lu et al. 1999). Therefore, even though ELT-2 NFs have likely experienced relaxed selection, they may still serve a function in these proteins, especially since they have not completely degraded. All but one of the *Caenorhabditis* species included in our study (the basal species, *C. plicata*) contain a singleton ELT-2 ortholog with an NF with sequence CX_2_CX_15-17_CX_2_C. Although most residues are non-canonical and not that well conserved, a few canonical GATA factor residues are conserved. These include a glycine (G) four residues upstream of the third ZnF-coordinating cysteine (found in 52 of 57 orthologs) (Fig. 3E; Supp. Fig. 13), which is highly conserved in non-*med Caenorhabditis* orthologs (Fig. 3A; Supp. Fig. 10,14) and canonical GATA factors (Teakle and Gilmartin 1998), and an acidic lysine (K) or arginine (R) at NF position 13 (found in 55 of 57 orthologs) (Fig. 3E; Supp. Fig. 12), which is a structurally important arginine in the canonical NF sequence that also directly interacts with bound DNA (Omichinski et al. 1993). Both the NFs of ELT-1 orthologs and their downstream basic regions are highly conserved when compared to canonical NFs of vertebrates (Fig. 3D) (Teakle and Gilmartin 1998; Lowry and Atchley 2000). In summary, the NFs of ELT-2 orthologs have diverged extensively from their ELT-1 counterparts and from each other, but why they have not been lost from more than a single species remains a mystery. Both the *C. elegans* ELT-1 and ELT-2 orthologs bind to canonical WGATAR DNA sites (Shim et al. 1995; McGhee et al. 2007; McGhee et al. 2009; Araya et al. 2014; Du et al. 2016; Wiesenfahrt et al. 2016). ELT-1 also binds non-canonical GATC DNA sites (Shim et al. 1995) and GATR followed by AGAT, 3 bps apart on the opposite strand (Araya et al. 2014) while ELT-2 has only been found to bind to single WGATAA sites (McGhee et al. 2007; McGhee et al. 2009; Du et al. 2016; Wiesenfahrt et al. 2016; Lancaster and McGhee 2020). This difference in binding preference between ELT-1 and ELT-2 orthologs could be due to the ELT-1 NF if it binds DNA and expands the ELT-1 binding repertoire.

##### *elt-1* and *elt-2* orthologs were not found on the same scaffold and are likely on different chromosomes in most Caenorhabditis species

We never found *elt-1* and *elt-2* orthologs on the same piece of assembled genomic DNA (Supp. Fig. 12). In each of the few species in this study for which there was chromosome level genome resolution available, the *elt-1* and *elt-2* orthologs were found on different chromosomes (4 and X, respectively). Through our synteny analysis (Fig. 4, see Materials and Methods) we predict that, with few exceptions, *elt-*2 orthologs lie on the X chromosome throughout the clade and *elt-*1 orthologs are found on chromosome 4.

#### elt3 clade

The topology of this clade, the origin of *elt-7*, *end-1*, and *end-3*, and how this elt3 clade expansion re-wired the endoderm specification network will be discussed elsewhere (Darragh AC, Rifkin SA, unpublished data, https://doi.org/10.1101/2022.05.20.492851, last accessed May 23, 2022). Here we only include a comparison of the GATA DBDs of these proteins.

##### elt3 clade GATA DBDs have residues that set them apart from other GATA DBDs

Representations of the consensus GATA DBDs for each elt3 clade ortholog group, created from pHMMs (Eddy 1998) of each, are shown in Figure 3A. Each of the elt3 clade pHMMs generally score the other clade proteins higher than other GATA proteins (Fig. 3B). Conserved residues within the elt3 clade GATA DBDs further support common ancestry among the ELT-7, END-1, END-3, and ELT-3 orthologs. For example, there is a glutamic acid (E) at position 17 in the ZnF loops of all confident representative ELT-7, END-1, END-3, and ELT-3 homologs (Fig. 3A; Supp. Fig. 14; see Materials and Methods; Supp. Fig. 16), and this residue is not conserved in any other canonical GATA factor groups (Supp. Fig. 10,13.15). (This residue is conserved in all singleton RCOR-1 ZnFs and in all but one singleton SPR-1 ZnFs, even though rcor1 clade ZnFs are quite divergent from canonical GATA factor ZnF motifs overall). (Supp. Fig. 3; Fig. 3A). An NMR structure of the DBD of cGATA-1 bound to DNA showed that the valine (V) at position 17 in the cGATA-1 ZnF loop is important for the structural integrity of the ZnF motif (Omichinski et al. 1993), and most non-MED canonical *Caenorhabditis* GATA factors also contain a valine at this position (Supp. Fig. 10,13).

Another residue uniquely found in all representative ELT-7 and ELT-3 homologs, and all but one END-3 ortholog, is an asparagine (N) serving as the first residue of the basic region (Supp. Fig. 14). This asparagine with its larger polar side chain is quite different from the small flexible non-polar glycine (G) residue conserved at this position in most canonical GATA factors (Supp. Fig. 10,13.15). Most END-1 orthologs have a serine (S), which is polar and smaller than asparagine but larger than glycine, at this position (Supp. Fig. 14). Two additional, adjacent residues are conserved in all representative ELT-3 homolog ZnFs and in most representative END-3 and END-1 homolog ZnFs: an arginine (R) followed by a glutamic acid (E) at positions three and four in the ZnF loop (Supp. Fig. 14). Arginine is also found in the same position in some rcor1 clade orthologs, and glutamic acid is also found in the same position in many MED orthologs and in some EGL-27 orthologs (Supp. Fig. 3,15,7). However, this combination of a negatively charged residue adjacent to a positively charged residue in these positions is not found in the ZnF motifs of any other proteins included in this study (Supp. Fig. 3,7,10,13,15), nor is it typical for other GATA factors (Teakle and Gilmartin 1998). These non-canonical residues that are conserved among the elt3 clade GATA DBDs support the monophyly of this clade.

##### ELT-3 and ELT-7 orthologs have more canonical animal GATA factor residues in their GATA DBDs than END-1 and END-3 orthologs

Figure 3C shows that the ELT-3 pHMM has 16 residues identical to and three residues similar to the 24 residues important for animal GATA factor DNA interactions and 12 of the 18 structurally important residues plus two similar residues (Omichinski et al. 1993; Bates et al. 2008; Lowry et al. 2009; Chen et al. 2012). The ELT-7 pHMM has 14 of the 24 residues important for DNA interactions plus four similar residues and 13 residues identical to the 18 structurally important residues (Fig. 3C). The END-1 pHMM has 13 residues identical to and three residues similar to the 24 DNA-interacting residues and 10 of the 18 structurally important residues plus two similar residues (Fig. 3C). The END-3 pHMM has 11 of the 24 DNA-interacting residues plus three similar residues and 11 residues identical to and one residue similar to the 18 structurally important residues (Fig. 3C). Overall, the ELT-3 and ELT-7 pHMMs have about the same number of conserved important residues (Fig. 3C), which may contribute to the similar binding preference of the *C. elegans* orthologs for TGATAA DNA sites (Gerstein et al. 2010; Narasimhan et al. 2015). The END-1 and END-3 pHMMs also have a similar number of conserved functional residues. However, they have fewer than the ELT-3 and ELT-7 pHMMs (Fig. 3C), which may be why *C. elegans* END-1 and END-3 have a similar binding preference of GATA DNA sites, but no or not as much specificity for flanking sequences, respectively (Narasimhan et al. 2015). The conservation of residues important for DBD binding and structure suggests that END-1 and END-3 orthologs have lost some of the residues that we presume they initially acquired from an ancestral *elt-3* or *elt-7* duplication. If this is the case, the *end* ancestor probably experienced weaker selection compared to its paralog and/or was selected for a broader range of binding.

#### MED ortholog group

More details about this ortholog group will be discussed elsewhere (Darragh AC, Rifkin SA, unpublished data, https://doi.org/10.1101/2022.05.20.492851, last accessed May 23, 2022). Here we only include an analysis of their GATA DBDs compared to that of other *Caenorhabditis* GATA-domain-containing proteins.

##### MED orthologs share non-canonical residues and have more variability in their GATA domains compared to other canonical *Caenorhabditis* GATA factors

We created pHMMs (Eddy 1998) of the *Japonica* group and *Elegans* group MED DBDs, respectively (Fig. 3A) and used them to query all the proteins in our analysis (Fig. 3B; see Materials and Methods). Except in ZnF loop length, *Japonica* group and *Elegans* group MED DBDs are more like each other than to any of the other GATA domains in this study (Fig. 3A) and so, as expected, their pHMMs scored each of the MEDs from the other group second highest, after the MEDs from their own species group (Fig. 3B). Both MED pHMMs specifically score MEDs higher than other GATA domain containing proteins (Fig. 3B), supporting the MED ortholog group in our phylogeny (Fig. 1A). Some primarily MED-specific residues are conserved within most MED DBDs, supporting predictions of their more recent common ancestry and divergent binding specificity, while other poorly conserved residues suggest these genes are experiencing rapid evolution and/or relaxed selection. For example, one of the similar residues arginine (R), lysine (K), or glutamine (Q) (Henikoff and Henikoff 1992) is usually found at position nine in MED ZnFs whereas similar residues are not found at this position in any of the other GATA domains (Fig. 3A). The arginine at this position in *C. elegans* MED-1 forms hydrogen bonds with the 5’ region of the non-canonical MED DNA binding site (Lowry et al. 2009), thus part of this non-canonical binding may be conserved in most, if not all, MEDs. On the other hand, a tyrosine (Y) found at position 19 in the BRs of most *Elegans* group MEDs is not conserved in *Japonica* group MEDs (Fig. 3A; Supp. Fig. 15) and in *C. elegans* MED-1 this tyrosine interacts with the 3’ region of the non-canonical MED DNA binding site (Lowry et al. 2009). Since this position was not found to be involved in the binding of other animal GATA factors to DNA, *Japonica* group MEDs may have a more canonical DNA binding site than *Elegans* group MED orthologs. Some poorly conserved residues in MED DBDs include position nine in the ZnF, which is a highly conserved threonine (T) in canonical GATA factors but a threonine, serine (S), asparagine (N), or cysteine (C) in MED orthologs (Fig. 3A; Supp. Fig. 15) and four residues before the third zinc coordinating cysteine which is a glycine (G) in most canonical GATA factors, but an asparagine, aspartic acid (D), glutamic acid (E), threonine, lysine, arginine, or serine in the MEDs.

Most (86 of 94) *Elegans* group MED ZnFs have loops 18 residues long (Fig. 3A; Supp. Fig. 4; Supp. Fig. 15). This is atypical for animal GATA factors and for the other genes in this study, although canonical plant and fungi GATA factors have loops of this length (Teakle and Gilmartin 1998). The eight other *Elegans* group MEDs all have ZnF loops that are 17 residues long (Supp. Fig. 4; Supp. Fig. 15), like canonical animal GATA factors (Teakle and Gilmartin 1998). These MEDs are found in three non-basal species, two of which are sister species. Due to the placement of the two sister species MEDs in our phylogeny (Fig. 1) the most parsimonious explanation is that a three nucleotide deletion, or an alternative splice site, in the ZnF loop coding sequence of one ancestral MED occurred in the most recent common ancestor of *C*. sp. *48* and *C. brenneri*. A *C*. sp. *51* MED-like protein also only has a 17 residue long loop, but this may be due to this protein having a more recent common ancestor with a non-MED GATA-domain-containing protein (see above). We think that the *Elegans* group ancestral MED had a ZnF with 18 residues in its loop. The ZnFs loops of most (23 of 36) *Japonica* group MEDs are also an atypical length - they are a residue shorter than canonical GATA factors (Supp. Fig. 4; Supp. Fig. 15) (Lowry and Atchley 2000) but the same length as most *Caenorhabditis* EGL-27 ZnF loops (Fig. 3A; Supp. Fig. 4; Supp. Fig. 15). However, due to the lack of homology between *Japonica* group MEDs and EGL-27 orthologs, this likely reflects convergent evolution. The other 13 *Japonica* group MEDs have ZnF loops of length 17 (Supp. Fig. 4; Supp. Fig. 15), like canonical animal GATA factors (Teakle and Gilmartin 1998), but since they are not found in basal *Japonica* group species, we propose that the ancestral *Japonica* group MED had a 16 residue long ZnF loop. Due to their placement in our phylogeny (Fig. 1) and since GATA ZnF insertions are relatively rare compared to substitutions, at least eight of these 13 *Japonica* group MEDs likely shared the same three nucleotide insertion, or alternative splice site, in their most recent common ancestor.

We compared the residue with the highest probability at each position in the *Japonica* group and *Elegans* group pHMMs (Fig. 3A) to the residues known to be important structurally or for DNA binding in animal GATA factors bound to DNA (Fig. 3C) (Omichinski et al. 1993; Bates et al. 2008; Lowry et al. 2009; Chen et al. 2012). Both of these MED pHMMs have nine of the 24 important DNA interaction residues, and four residues that are similar. Of the 18 structurally important residues, the *Japonica* group and *Elegans* group pHMMs have eight and 10 identical and three and two similar residues, respectively (Fig. 3C). Compared to the EGL-27 and rcor1 clades, MED DBDs contain a couple more residues important for GATA factor DNA binding or structural integrity in common with canonical animal GATA domains (Fig. 3C). Moreover, MED GATA domains more closely resemble canonical *Caenorhabditis* GATA domains than they do the atypical EGL-27 or rcor1 clade GATA domains (Fig. 3A), supporting the hypothesis that the MEDs arose from one of the canonical *Caenorhabditis* GATA factors instead of from a different GATA-domain-containing protein.

## Discussion

### Radiation of GATA-domain-containing proteins around the time of the *Elegans* supergroup ancestor

We found additional evidence for Maduro’s (2020) proposed expansion of GATA factors – *end-1*, *end-3*, *elt-7*, and *med* – in the ancestor of the *Elegans* supergroup and predict that a duplication of the *Elegans* supergroup ancestor *rcor-1* produced the *rcor-1* and *spr-1* orthologs found in contemporary species (Fig. 1; see Results; Darragh AC, Rifkin SA, unpublished data, https://doi.org/10.1101/2022.05.20.492851, last accessed May 23, 2022). Moreover, we found that a tandem *elt-6* duplication likely produced *egl-18* in the *Elegans* supergroup/*Guadeloupensis* group ancestor (Fig. 1; see Results). It is intriguing that these six duplications of GATA-domain-containing regulatory proteins likely occurred around the same time, and it will be interesting to look genome wide to see whether other protein families also expanded (or contracted) at this time, particularly those involved in development. There is a lot of variation in protein-coding gene count between the 48 likely non-heterozygous *Caenorhabditis* draft genomes analyzed here (Stevens 2020), suggesting that protein-coding genes duplication and losses are frequent across the genus. Moreover, the finding that the spontaneous gene duplication rate in *C. elegans* was three orders of magnitude larger (Konrad et al. 2018) than the point mutation per nucleotide site rate (Denver et al. 2009), suggests that gene duplication would be a quicker way to increase the gene expression of a locus than nucleotide substitution (Lipinski et al. 2011). In fact, evidence in multiple natural populations support this idea (Nair et al. 2007; Perry et al. 2007). Not only did this radiation re-wire at least the endoderm specification network (Maduro 2020) but it is associated with changes in selection pressures on at least one paralog (Fig. 5,6; Darragh AC, Rifkin SA, unpublished data, https://doi.org/10.1101/2022.05.20.492851, last accessed May 23, 2022). These duplications are not associated with any known change in the environment or morphology of these animals and so the evolutionary forces driving this change remain unknown.

**Figure 6.**
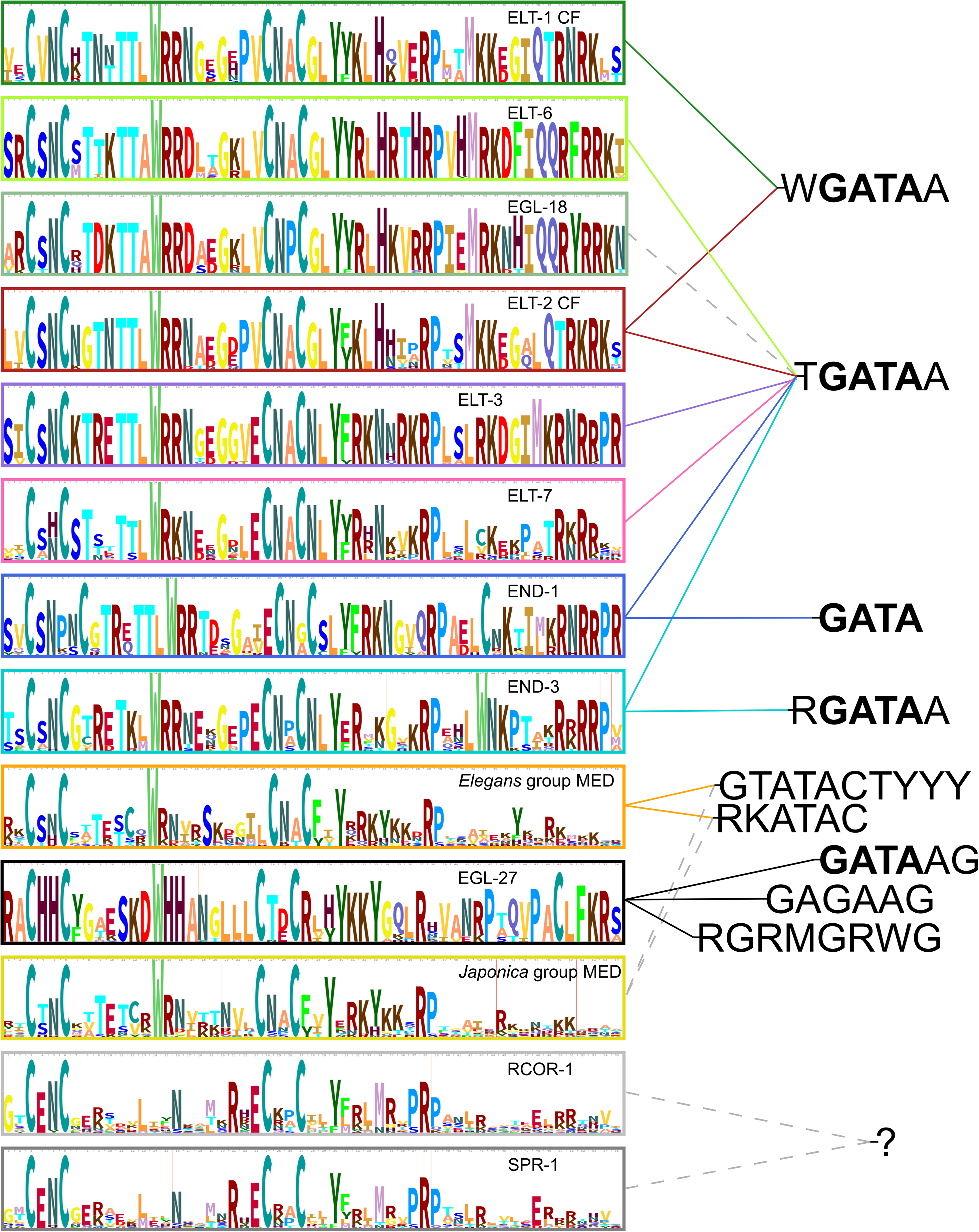
Diversification of *Caenorhabditis* GATA-like DNA-binding domain sequences and their DNA binding targets in *C. elegans*. CF/ZnF pHMMs representing the 13 GATA DBD groups are shown on the left, ordered by their number of conserved non-MED animal GATA factor structure and DNA interaction residues (those at the top with the most and those at the bottom with the least). DNA sequences that the *C. elegans* orthologs of these groups bind to are shown on the right; the solid colored lines from the pHMMs to the DNA sequences indicate binding interactions supported by experimental evidence from the literature; dashed gray lines indicate unknown and, in the case of the *Japonica* group MEDs, hypothesized binding interactions. The DBD group colors are the same as used in Figure 3.

### Functional divergence under similar selection strengths

After a full-length gene duplication (including regulatory regions) presumably paralogs would be completely redundant and therefore shielded from selection as long as the ancestral function is maintained through their combined abilities (Ohno 1970). Through this process of relaxed selection, paralogs would diverge rather quickly, that is until they have non-redundant functions that would be selectively maintained (Sidow 1996). We did not find significant relaxation of selection pressures on *spr-1* orthologs compared to *Elegans* supergroup *rcor-1* orthologs nor between *Elegans* supergroup and non-*Elegans* supergroup *rcor-1* orthologs (Fig. 5). This is consistent with the rcor1 branch lengths, which are similar for these groups (Fig. 5). This suggests that the strength of selection didn’t change significantly after duplication even though the *C. elegans* SPR-1 and RCOR-1 orthologs have non-redundant functions (Jarriault and Greenwald 2002; Bender et al. 2007; Hale et al. 2014; Vandamme et al. 2015). Therefore, the divergence of these paralogs likely occurred under a similar selection strength as their singleton ancestor. It would be interesting to uncover the function of non-Elegans supergroup rcor-1 orthologs to see if this divergence involved subfunctionalization, neofunctionalization, or a combination.

### Relaxed selection on both newer paralogs

We did not find evidence for significant relaxed selection on the elt6 clade paralogs relative to each other (*egl-18* orthologs relative to the *Elegans* supergroup and *Guadeloupensis elt-6* orthologs) relative to each other (Fig. 5). *C. elegans egl-18* and *elt-6* have overlapping functions and are even sometimes transcribed as a dicistron, therefore they seem to have been maintained for the same role. Having similar selection pressures is consistent with this. We did detect relaxation of selection on *Elegans* supergroup and *Guadeloupensis elt-6* orthologs compared to the other *Caenorhabditis elt-6* orthologs (Fig. 5). This is consistent with the ELT-6 ortholog group branch lengths, which have different lengths for these groups (Fig. 5). This suggests that after duplication both the *elt-6* and *egl-18* paralogs have experienced relaxed selection compared to their ancestral singleton *elt-6*. Perhaps this relaxation of selection has led to changes in function of these homologs.

### GATA domain intron/exon structure supports the clades in our phylogeny but is not sufficient to resolve the complete *Caenorhabditis* GATA factor birth order

Recent non-retrotransposed gene duplicates will often share gene structure characteristics, such as locations of introns and domains, which can help untangle the evolutionary history of a gene family (Gillis et al. 2008; Eurmsirilerd and Maduro 2020; Maduro 2020). Maduro (2020) found conservation of an intron in the coding sequence for the ZnF of all *end-1*, *end-3*, and *Japonica* group *med* homologs he examined and loss of this intron in *Elegans* group *med* homologs, which mostly lack introns completely. The same intron location, immediately before the last seven nucleotides that code for the ZnF, was also found in *C. elegans elt-2* CF (Fukushige et al. 1998) and *elt-7* (Sommermann et al. 2010) and some additional (but not specified) *Elegans* supergroup species *elt-2* and *elt-7* orthologs (Maduro 2020). In addition, this splice site location was found in most nematode *elt-2* and *elt-3* orthologs examined, including five *Caenorhabditis* species (Eurmsirilerd and Maduro 2020). However, this differs from most nematode Clade I *elt-2* orthologs which instead have a CF intron at the same location as most nematode *elt-1* CFs, the *Drosophila melanogaster* (fruit fly) GATA factor *Grain* CF, and the cGATA-1 CF (Eurmsirilerd and Maduro 2020), supporting the elt1/2 clade found in our phylogeny (Fig. 1A). We found an intron immediately before the last seven base pairs that code for the ZnF in most of the additional *end-1, end-3*, *elt-2*, *elt-3*, and *Japonica* group *med* orthologs included in this study (Fig. 2; Darragh AC, Rifkin SA, unpublished data, https://doi.org/10.1101/2022.05.20.492851, last accessed May 23, 2022), corroborating the findings of both Maduro (2020) and Eurmsirilerd & Maduro (2020). Only one *C*. sp. *51 end-3* paralog, two *C. macrosperma med* paralogs, and three *elt-2* orthologs have lost this intron (Supp. Fig. 2). Moreover, we found an intron at this location in all *elt-7*s (Supp. Fig. 2). This shared intron between *end-1*, *end-3*, *elt-2*, *elt-3*, *elt-7*, and *Japonica* group *med* orthologs suggests common ancestry. More recent shared ancestry between *end-1*, *end-3*, *elt-3*, and *elt-7* is captured by our phylogenetic analysis (Fig. 1), our GATA domain pHMMs (Fig. 3A), and the pHMM scores (Fig. 3B), but none of these methods provide evidence of additional homology for *elt-2* and *Japonica* group *med* orthologs. Extensive sequence divergence may have obscured any evolutionary relationship.

We did not find this conserved splicing location in any of the other GATA domain-containing proteins in our study. However, we did find other conserved intron locations within and between clades. In corroboration with previous results (Eurmsirilerd and Maduro 2020) we found conservation of the *elt-1* CF intron location in the basic regions of all singleton confident *elt-6s*, *egl-18*s, and *elt-1* CFs (Supp. Fig. 2). This shared intron supports the more recent common ancestry of *elt-6* and *egl-18* orthologs in our phylogeny (Fig. 1). However, more distant evolutionary history is still not clear. This is because the ancestral elt6 clade GATA factor likely originated at the base of Chromadoria, and the only two GATA factor orthologs in Clade I nematodes, *elt-1* and *elt-2* orthologs, both had an intron conserved at the same position in their CF BRs at this time (Eurmsirilerd and Maduro 2020). Thus, it is unclear whether an *elt-1* or an *elt-2* duplication produced the elt6 clade ancestor (Eurmsirilerd and Maduro 2020). EGL-27 and rcor1 clade orthologs also have conserved but different splice sites in their GATA-like domains respectively (Supp. Fig. 2), suggesting that these domains have been diverged from canonical GATA factors at least since the *Caenorhabditis* ancestor.

### *Caenorhabditis* GATA domain diversification, while likely maintaining a binding preference for GATA DNA sites

For the most part our GATA-domain-only phylogeny (ZnF tree) has the same 12 ortholog groups as our full-length tree (Supplemental Figure 5 compared to Figure 1). However, the topologies of these trees differ and the ZnF tree contains the elt1/2 clade NFs and CFs. For example, some of the larger clades and ortholog groups found in the full-length tree are not monophyletic in the ZnF tree (i.e., elt1/2 clade CFs, elt3 clade ZnFs, ELT-1 NFs and CFs, EGL-18 ZnFs, ELT-7 ZnFs, END-1 ZnFs, SPR-1 ZnFs, and MED ZnFs) and the other clades are less well defined. This is likely primarily because the ZnF tree does not contain as much information about the evolutionary history of these proteins: it still captures the uniqueness of the 14 ZnF groups but some of the other relatedness between the groups has been lost. Most orthologous *Caenorhabditis* GATA domains lack sequence variability and have likely experienced negative selection (Fig. 3A) which can also be visualized by the short branches within most ortholog groups in the ZnF tree (Supp. Fig. 5). However, there are a few exceptions. The *elt-2* NFs, are highly variable, lack many highly conserved canonical GATA domain residues (see Results; Fig. 3E; Supp. Fig. 13), and are on divergent long branches (Supp. Fig. 5). The RCOR-1 and SPR-1 GATA-like domains lack some highly conserved canonical GATA domain residues (see Results; Fig. 3A; Supp. Fig. 3) and are more variable than all non-MED *Caenorhabditis* GATA domains (Supp. Fig. 5). The MED GATA domains are the next most variable, but this is more expected due to the quick turnover of this group (Supp. Fig. 5; Fig. 3A; Supp. Fig. 15; Fig. 1A) (Maduro 2020). The *C. elegans* orthologs from all other ortholog groups besides EGL-18 which has not been tested and including ELT-2s CF bind to canonical GATAA sites (Shim et al. 1995; McGhee et al. 2007; McGhee et al. 2009; Xu and Kim 2012; Araya et al. 2014; Narasimhan et al. 2015; Du et al. 2016; Wiesenfahrt et al. 2016), suggesting that even though these GATA domains are divergent they still bind to similar DNA sequences (Fig. 6). Overall, GATA DBDs appear to occupy a large sequence space, but possibly a restricted functional space. It will be interesting to investigate whether the similar DNA binding is primarily due to the similar structures of these domains or if there is flexibility in the structure and residues that bind to similar GATA sequences. Additionally, any binding differences, like that of *C. elegans* MEDs, could be due to non-DBD sequences, which was found to be the predominant case for whole-genome duplicate transcription factors in *Saccharomyces cerevisiae* (yeast) (Gera et al. 2022).

### *Caenorhabditis* GATA domain hidden Markov model profiles can be used to help identify orthologs in newly sequenced *Caenorhabditis* and other nematodes

We created pHMMs (Eddy 1998) representing the GATA DNA-binding domains of the 12 ortholog groups in our phylogeny (Fig. 3A). We searched for alignments of these pHMMs against most proteins included in this study and found that even these relatively short profiles could clearly distinguish between the 13 DBD groups (Fig. 3B). Moreover, the profiles of ortholog groups in larger clades in our tree, scored their adjacent ortholog group(s) next highest (e.g., EGL-18 profile scored ELT-6 profile second highest) (Fig. 3B), suggesting that at least some of the phylogenetic signal is captured in the 55 residues of these profiles. Additionally, we used these pHMMs to score paralogs that were not used to create the profiles. Similar to singletons, paralogs scored highest on their ortholog groups pHMM (data not shown). The inclusion of up to 58 *Caenorhabditis* species GATA domain sequences to create these pHMMs, the comparable speeds of HMMER searches to those of BLAST, and the specificity of these profiles suggests that they can be used to identify orthologs and even divergent paralogs in newly sequenced *Caenorhabditis* species and likely other nematode species as well. Our search for GATA factor orthologs used the PROSITE GATA-type zinc finger domain profile, which was designed to detect distantly related GATA domain-containing proteins, which is why in addition to canonical GATA factor domains from fungi, plants, and animals this profile included *C. elegans* EGL-27 and SPR-1 atypical GATA domains (prosite.expasy.org). In addition to the *Caenorhabditis* GATA factor ortholog group specific profiles (Fig. 3A) we created a more general *Caenorhabditis* GATA factor profile that included ELT-6, EGL-18, ELT-1 CF, ELT-2 CF, ELT-3, ELT-7, END-1, END-3, and MED GATA domain sequences in its creation (data not shown). We hope that these *Caenorhabditis* GATA factor pHMMs will be a valuable resource to the *Caenorhabditis* and greater worm community.

### Neofunctionalization of at least the *Elegans* group *med* ancestor for binding to a non-GATA DNA site and intercalation into the mesoderm specification network

The *C. elegans meds* and at least some *C. briggsae* and *C. remanei meds*, function early in both mesoderm and endoderm specification networks (Maduro et al. 2001; Coroian et al. 2006). The *C. elegans meds* bind to a non-GATA DNA sequence (Broitman-Maduro et al. 2005; Lowry et al. 2009). Since some *C. briggsae* and *C. remanei meds* are able to compensate for lack of both the *C. elegans meds*, it is likely that they similarly bind GTATACTYYY instead of canonical HGATAR sites. This non-canonical binding and function in the mesoderm is unique among *C. elegans* GATA factors (Hawkins and McGhee 1995; Shim et al. 1995; Zhu et al. 1997; Gilleard et al. 1999; Gilleard and McGhee 2001; Koh and Rothman 2001; Maduro and Rothman 2002; Fukushige et al. 2003; Narasimhan et al. 2015; Du et al. 2016; Wiesenfahrt et al. 2016). Thus, these novel features likely arose in an ancestral *med*. To narrow down the timeline, it would be fruitful to examine whether *Japonica* group *med* expression, binding, and function mirror that of *Elegans* group *meds* or that of canonical GATA factors. If these novel properties are conserved, it would suggest that the ancestral *med* neofunctionalization occurred in the *Elegans* supergroup ancestor. If not, we would predict that they arose in the *Elegans* group ancestor, unless the binding data from *C. elegans* is truly unrepresentative.

### Another example of developmental system drift in *Caenorhabditis*

Other than male tails, *Caenorhabditis* species are very similar anatomically (Kiontke et al. 2011; Slos et al. 2017) and are thought to have similar cell lineages as *C. elegans*, since the lineages of *C. briggsae* (Zhao et al. 2008) and *Pristionchus pacificus* (Vangestel et al. 2008) are almost identical to that of *C. elegans* (Sulston et al. 1983). Therefore, we think that the GATA-domain-containing duplications, redundancy, and neofunctionalization that we have described here and that have changed developmental networks have had little overt effect on the developmental output of these animals. This phenomenon has been termed developmental system drift (True and Haag 2001) and has been previously documented in *Caenorhabditis* (Félix 2007; Ellis and Lin 2014; Verster et al. 2014; Haag et al. 2018; Maduro 2020). We expect that additional examples of developmental system drift in *Caenorhabditis* will continue to be uncovered as development in more species and strains is characterized in greater detail.

## Conclusion

Developmental genes are not necessarily evolutionarily constrained even if development is. They can be duplicated and diverge or be maintained as redundant paralogs, all without drastic changes to animals’ anatomy.

## Materials and Methods

### Identifying GATA factor homologs

We downloaded the proteome files for 56 *Caenorhabditis* species and two *Diploscapter* species, and the transcriptome files for *C.* sp. *45* and *C*. sp. *47* from Caenorhabditis.org (the *Caenorhabditis* Genome Project) in early 2020. We searched for GATA DNA-binding domains in the proteome and transcriptome files that matched the PROSITE GATA-type ZnF domain profile PS50114 (prosite.expasy.org) using the pftools3 (https://github.com/sib-swiss/pftools3) pfsearchV3 tool (Schuepbach et al. 2013). We identified 890 proteins which had at least one match of score eight or more and used many of them (and a few more, see below) for this study (while the rest will be reported on elsewhere).

For any case in which we expected to identify an ortholog of a *C. elegans* GATA-domain-containing protein but did not, we performed reciprocal protein-protein BLAST (BLASTp) searches using the Biopython NcbiblastpCommandline wrapper and/or protein query-translated subject BLAST (tBLASTn) searches using the NcbitblastnCommandline wrapper (Altschul et al. 1990; Camacho et al. 2009; Cock et al. 2009) against the *C. elegans* ortholog, and/or sister species ortholog(s), to identify any additional GATA-domain-containing proteins we may have missed. tBLASTn searching was performed on scaffold files for species of interest downloaded from Caenorhabditis.org (the *Caenorhabditis* Genome Project) in early 2020. We used an e-value cutoff of 0.001 to non-*C. elegans* species and an e-value cutoff of 0.1 back to *C. elegans* for these BLAST searches. We identified an additional 48 EGL-27, RCOR-1, or SPR-1 homologs (which will be reported on elsewhere), an ELT-7 homolog (in *C. becei*), a MED homolog (in *C. panamensis*, referred to as CPANA.med), and another MED homolog (in *C. macrosperma*, referred to as CMACR.med), using this method. We suspect that many of the EGL-27, RCOR-1, and SPR-1 homologs that we identified through BLAST searches were not identified using the PROSITE profile because their ZnF motifs are too divergent. On the other hand, the ELT-7 and MED homologs were identified using BLAST but not using PROSITE because they contained annotation errors; the annotation of the *C. becei* genome *elt-7* contained a premature stop codon in its ZnF, for example, and the two putative MED homologs in the genome sequences of *C. panamensis* and *C. macrosperma* were not annotated as having coding regions.

Although we may have missed additional proteins with GATA-like DNA-binding domains in our searches, any genes encoding them would likely be undergoing pseudogenization or on some dramatically different evolutionary trajectory than the sequences that we included in our analysis; it is also possible we might have missed a few other proteins due to additional annotation errors. Additionally, some of the paralogs we included in our study may represent artifacts related to possible heterozygosity in the sequenced strain(s) and the quality and coverage of the genome assemblies for some of the species (Barrière et al. 2009; Haag and Thomas 2015; Stevens 2020), this possibility being most likely for paralogs in the MED ortholog group which we often found contained many highly similar species-specific sequences (Fig. 1).

### Phylogenetic analysis

Phylogenetic analysis is dependent on alignment of multiple sequences, but sequence alignments can be noisy and dependent on arbitrary factors like sequence direction and the order of sequences in a list. Furthermore, when aligning many sequences, some will be alignable over greater lengths than others. This is particularly true in our case where all the sequences share a common alignable core GATA ZnF but can be otherwise so different that they fall into different subfamilies. Within a subfamily, sequences will be alignable over greater distances than between subfamilies, and this is evident in our alignments. A single alignment of multiple sequences, however, provides no way to estimate how variable the alignment between any two sequences is at any given residue and therefore no way to weight the information at each aligned position by sequence pair combination for downstream applications like phylogenetic inferences or population genetic estimations. To capture the variation in alignability, we followed an approach like that used by Penn and colleagues(Penn et al. 2010) and generated a heterogeneous set of alignments. The rationale is that alignable regions should be fairly impervious to variations in the guide tree used to build the alignment or to the sequence direction, while unalignable regions will be sensitive to these manipulations; thus. across a collection of multiple alignments, the consistent signal from alignable residues will contribute more than the conflicting noise from poorly aligned regions. Our multiple alignments were constructed following 5 steps: (1) divide the sequences into separate groups (see detailed description below for justification); (2) build an initial alignment; (3) bootstrap this alignment; (4) generate neighbor-joining (NJ) trees from these bootstrapped alignments; (5) serially align the sequences using these NJ trees as guide trees.

Only 884 of the total of 941 putative protein sequences we examined contained a well-aligning GATA-like DNA-binding domain. We aligned the longest isoforms of these 884 protein sequences using the default options of Clustal Omega (Sievers et al. 2011). By construction these sequences include a common core domain that varies slightly in residues and length; beyond that domain these proteins are much less conserved. Because Clustal Omega uses pHMM alignment (Sievers and Higgins 2018), it is ideal for aligning sequences like these and generated multiple alignments with excellently aligned GATA ZnFs that were much more compact than, for example, alignments generated using MAFFT (Katoh et al. 2002). However, when we aligned all 884 sequences at once, a few invariably were nearly completely offset from the rest. To overcome this, we serially aligned the sequences in batches. After the first batch of sequences was aligned, an pHMM of the alignment was used to align them with the next batch of sequences, and this process was repeated until all sequences were aligned. We also constrained the alignments to always align with the first cysteine in the CF. These procedures resulted in alignments of the conserved residues in the CF. END-1 orthologs have an extra two residues between the first pair of cysteines in their ZnFs; to create a consistent alignment, we therefore set the sequence of non-END-1 proteins to CX--XC.

After initially aligning the 884 sequences, we bootstrapped (Felsenstein 1985) these alignments and used them to make new NJ guide trees (Saitou and Nei 1987). We then used each new guide tree to make a new Clustal Omega (Sievers et al. 2011) alignment using the serial alignment approach (described above) for sequences in both (forward and reverse) directions. We repeated this process 10 times and then randomly chose nine of these alignments (five forward; four reverse) and concatenated them to the original, reversing the orientation of the reversed ones so that all sequences were in the same direction. This procedure effectively weights each position x sequence pair combination by how often they are consistently aligned. Using IQ-TREE 2 version 2.1.2 ModelFinder (Kalyaanamoorthy et al. 2017) on the Cyberinfrastructure for Phylogenetic Research (CIPRES) Science Gateway V. 3.3 (phylo.org) we identified the VT+F+I+G4 model (variable time (Müller and Vingron 2000), empirical base frequencies from supplied alignment, allowing invariable sites, and discrete Gamma model (Yang 1994) with four rate categories, respectively) as the best model of evolution for this alignment. We then used the IQ-TREE 2 version 2.1.2 tree inference tool (Minh et al. 2020) to estimate the evolutionary history of these sequences. The resulting maximum likelihood phylogeny of all 884 proteins is shown in Figure 1A.

### Process for editing gene annotations

Upon visual inspection and comparison of ortholog group sequences it became clear that some of the gene annotations in the files used for this study were likely incorrect because their coding sequences were highly divergent from the majority of their orthologs and considerably less divergent coding sequences were possible upon making only minor adjustments. We therefore judiciously edited some annotations to make the sequences more homologous. For example, the initial annotations of a few genes contained premature stop codons, but by simply adding an alternative splice site these coding sequences became full-length sequences with much better sequence homology to their orthologs. Additionally, we identified multiple examples of cases in which multiple genes had been annotated as a single gene, and parts of genes had been annotated as multiple genes, and addition of alternative splice sites to these genes resulted in improved homology among their orthologs as well. In total we slightly edited like this the annotations of 226 genes. We annotated coding sequences instead of exons because the data for a few of the 60 species include untranslated regions in their exon sequences; although most of the data files did not contain this information, we wanted consistent sequences that started from the first coding ATG for our analyses. Notes on the types of edits we made are included under the “editingNotes” column in Supplemental Table 1. These edits increased the number of genes we considered “confident” (see below), in that we were confident in using them for further studies and in the robustness of the phylogeny they were used to create.

### Identifying confident sequences for further analyses

For all additional analyses of the protein sequences included in our phylogeny we focused on a subset of the 884 that, based on their sequence features and how robustly they grouped into a clade in our preliminary phylogeny, best met important criteria for our studies. For most gene families, highly divergent protein sequences are likely on different evolutionary trajectories compared to their conserved relatives and may even be pseudogenizing; we therefore had less confidence in those highly divergent protein sequences and did not include them in further analyses. For example, we did not include protein sequences that were positioned on long individual branches at the base of an ortholog group, or comprised exceptionally long branches elsewhere, and/or had weak branch support values. The features of protein sequences deemed “not_confident” (see the “confidence” column in Supplelmental Table 1) can be found in either the “geneQuality” column and/or the “notes” column of Supp. Table 1. Proteins that grouped basally in a group are labeled “basal_group_name” (e.g., basal_med) in the “clade” and “orthologGroup**”** columns (Supp. Table 1).

To assess gene annotation quality, in early 2020 we downloaded from Caenorhabditis.org (the *Caenorhabditis* Genome Project) the annotation and scaffold files for the 56 *Caenorhabditis* species and two *Diploscapter* species for which we had obtained proteome files for this study and examined the protein-coding and neighboring sequences for each protein. Using a custom Python script, we extracted the coding sequence for each protein from that species’ annotation file, starting with the first ATG, if present (see “exonSeq” column in Supp. Table 1). We then marked whether the gene had features listed under the “geneQuality” column in Supplemental Table 1 that reduced our confidence in it. These features included: premature stop codons (“prematureStop”); lack of an obvious GATA ZnF (“noZnF”); fewer than 13 amino acid residues coded for after their ZnFs, suggesting an incomplete basic region (“shortBR”); lack of conservation/alignment in the sequences following the ZnF (“noBR”); absence of a start codon (“noMet”); absence of a stop codon (“noStop”); or truncation as compared to its orthologs (“truncatedStart” or “truncatedEnd”). There were also three protein sequences marked as either “nonHomologousBR”, “NsInGene”, or “NsUpsteamOfTruncation” which, as these labels imply, had either a non-homologous basic region, an incomplete sequence (i.e. unknown amino acids, AKA N’s) in the coding sequence, or N’s upstream of a putative gene truncation. If a gene’s annotation passed all our criteria, it was labeled “good” in the “geneQuality” column (Supp. Table 1).

We considered CX_2,4_CX_7_WX_9_CX_2_C as the canonical GATA ZnF motif for this study because this pattern, of similarly spaced four cysteines (C’s) and a tryptophan (W) at position 8 in the ZnF loop, is found in the CFs of all canonical animal GATA factor domains (Teakle and Gilmartin 1998; Lowry and Atchley 2000). In addition, to ensure that we did not miss any possibly homologous ZnFs, we included the following divergent ZnF motifs in our analysis: CX_2_CX_15-17_CX_2_C for ELT-2 ortholog N-terminal ZnFs (NFs), CX_2_CX_7_WX_8_CX_2_C for some *Japonica* group MEDs and most EGL-27 orthologs, CX_2_CX_16-17_CX_2_C for all RCOR-1 orthologs, and CX_2_CX_17-18,20-21,23_CX_2_C for all SPR-1 orthologs (ELT-2 NFs, EGL-27, RCOR-1, and SPR-1 results will be reported elsewhere). Protein sequences that lacked the expected GATA-like ZnF motif of the ortholog group that the protein grouped into are marked “not_confident” (see Supp. Table 1). Of the 31 protein sequences that were added to our analysis based on reciprocal BLASTp searches against orthologs in other species because no significant matches to the PROSITE GATA-type ZnF domain had been found for them among the proteome files (see above), 16 did not have an obvious GATA-like ZnF (“noZnF”) as a result of this analysis either, as expected. Most of the remaining 15 “noZnF” protein sequences appeared to have degrading/non-functional ZnFs since they contained only two or three of the usual four zinc-coordinating cysteines or, for those that had four ZnF cysteines, had fewer than 15 residues between the two cysteine pairs, a pattern which is not found in any canonical GATA factor ZnFs. MED sequences that had multiple ZnFs were also considered “not_confident” (see Supp. Table 1) because most MEDs (and all non-ELT-1/-2 *Caenorhabditis* GATA factors orthologs) have only a single ZnF (Lowry and Atchley 2000; Eurmsirilerd and Maduro 2020; Maduro 2020).

Overall, proteins that grouped robustly into a clade in our phylogeny, were classified as “good” in the “geneQuality” column in Supplemental Table 1, and contained an expected GATA(-like) ZnF motif for the ortholog group that the protein clustered into, were deemed “confident”. A total of 714 of the 884 proteins with well-aligning GATA-domains fit this description and were used to create the phylogeny shown in Figure 1B.

### Clade and ortholog group terminology

We refer to the 12 ortholog groups revealed in our phylogenies (Fig. 1) by the name of the *C. elegans* protein(s) within that group. For example, the group containing the *C. elegans* ELT-3 ortholog, and all the proteins from the other 59 species that we classified as ELT-3 orthologs, is referred to as the ELT-3 ortholog group (Fig. 1). We gave the ortholog groups that grouped adjacent to another ortholog group(s), thus forming larger clades, the name of the most ancient ortholog group within that larger clade, formatted in lower case, without a hyphen. For example, we refer to the larger clade containing the ELT-3, ELT-7, END-1, and END-3 ortholog groups as the elt3 clade (Fig. 1). However, one clade contains two ortholog groups that both contain orthologs from all species included in this study, so their evolutionary history is beyond the scope of our analysis, and we refer to this clade as elt1/2 to represent both the ancient ELT-1 and ELT-2 ortholog groups within it (Fig. 1).

### Gene structure comparisons and predictions of ancestral gene structures

Using a custom Python script, the exon sequences listed in the Supp. Table 1 column “exonSeq” (see above), and the respective scaffold sequence we determined intron lengths in our selected genes (data not shown). Using the ELM2 (PF01448), BAH (PF01426), and Myb (PF00249) Pfam domain seed alignments (pfam.xfam.org) and HMMER v3.3 tools (hmmer.org), we made pHMMs of each of these domains. Then we used HMMER v3.3 tools to search for pHMM domain matches (with no significance cutoff, default settings) in each protein sequence and found its corresponding location in each protein’s gene structure. Using a custom Python script we found the locations of the GATA ZnF domains that we identified in each confident protein (Supp. Fig. 4; see above). Using the exon lengths and the domain location information, we created representations of the gene structures of all the confident genes in this study for which genomic data was available (Supp. Fig. 2) using a custom R script. (Note: the *C*. sp. *45* and *C*. sp. *47* genes were excluded because only transcriptome data were available for these species.)

We visually compared the gene structures of 714 confident genes (Supp. Fig. 2) and, using the principle of parsimony (and when parsimony was not sufficient to distinguish between two alternatives also treating intron loss as more frequent than intron gain (Roy and Penny 2006)), then predicted some ancestral gene structures (exon number and domain location(s)) for each ortholog group (Supp. Fig. 2; Fig. 2). To estimate the lengths of the exons and introns in the ancestral genes, we calculated and used the median lengths of the exons and introns of the orthologs that had the same gene structure as the predicted ancestor (Supp. Fig. 2).

### Construction, comparison, and use of hidden Markov model profiles for ortholog group

We aligned singleton protein sequences (see above) within each of the ELT-1, ELT-2, ELT-6, EGL-18, RCOR-1, SPR-1, and EGL-27 ortholog groups, the singleton and representative paralog (see above; Supp. Fig. 16) protein sequences within each of the ELT-3, ELT-7, END-1, and END-3 ortholog groups, and all *Elegans* group and *Japonica* group protein sequences within the MED ortholog group respectively, using MUSCLE (Edgar 2004), MAFFT FFT-NS-2 (Katoh et al. 2002), and Clustal Omega (Sievers et al. 2011) default settings. Additionally, we aligned all protein sequences that we considered GATA factors (i.e. those in the elt1/2, elt6, and elt3 clades and in the MEDs ortholog group). Overall, all three alignment algorithms aligned the ZnFs similarly, although the surrounding regions contained more differences. Upon visual inspection, we concluded that the MUSCLE alignments did the best job of aligning the most ZnF- neighboring residues (i.e. introduced more gaps than Clustal Omega, but fewer than MAFFT, such that likely conserved residues were aligned between orthologs) and, therefore, MUSCLE was used for all alignments with the exceptions of the ELT-2 NFs and the all-GATA-factor alignments for which MAFFT and Clustal Omega, respectively, were used instead.

We trimmed the MUSCLE alignments to three different sizes. The small alignment includes the GATA ZnF and part of the adjacent BR up to the well conserved arginine (R), proline (P) pair which is usually located 12 to 13 residues after the end of the ZnF (Lowry and Atchley 2000; Eurmsirilerd and Maduro 2020; Maduro 2020). The medium-sized alignment includes the two residues before the start of the GATA ZnF and most, often all, of the adjacent BR (i.e. 28 residues after the ZnF), which contains all the residues involved in the structure and DNA-binding of the cGATA-1 CF (Omichinski et al. 1993)(Omichinski et al. 1993) and *C. elegans* MED-1 (Lowry et al. 2009). The large alignments comprise all reasonable-looking alignment positions surrounding the ZnF on either side, but not including any positions from non-GATA DNA-binding domains, as determined by visual inspection. Each category of truncated alignment was used to make two pHMMs using the HMMER version 3.3 hmmbuild tool (hmmer.org), one with default and the other with enone settings. (Enone uses the actual number of aligned sequences for the effective number of sequences, which maximizes the information content per position.) We then used hmmscan with no bit score cutoff (Mistry et al. 2013) to identify and score profile matches among all the protein sequences included in this study. Enone medium pHMMs are shown and used in Figure 3. We used a custom Python script to create radar plots depicting the bit scores for the protein sequences (see Fig. 3B). We used Skylign and its “information content above background amino acid frequencies” option (Wheeler et al. 2014) to make logos of the pHMMs (Fig. 3A,D,E). This option displays the total information content per position as the total height of the stack of amino acid(s). Only amino acid(s) with frequencies at that position above the background frequency of that amino acid in the BLOSUM62 substitution matrix (Henikoff and Henikoff 1992) are included in the stack.

### pHMM comparison to residues known to be important for animal GATA factors bound to DNA

We compared the residue with the highest probability at each position in each pHMM to the residues known to be important for the protein structure and/or DNA binding of animal GATA factors bound to DNA. We considered residues similar if they have a BLOSUM62 substitution score of 1 or higher (Henikoff and Henikoff 1992). The one exception was substitutions of lysines (K) with glutamic acid (E), in the basic region of the GATA domain. This exception was only implemented once for the MEDs which have a glutamic acid instead of a lysine as position 17 in their basic regions.

### Identifying syntenic GATA-domain-containing genes

Using a custom Python script, we analyzed the annotation files for each species to identify syntenic genes. We thereby established the scaffold/chromosome containing the sequence coding for each of the proteins in this study, and then determined whether any of those sequences were on the same scaffolds. The syntenic GATA-domain-containing genes, their shared scaffold, and the distance between these syntenic genes are listed in Supplemental Table 3. Since most species genome assemblies lack chromosome-level resolution we also used a custom Python script, to find all annotated genes within 70 kb upstream and downstream of each confident gene (“neighbor genes”). We then used BLASTp (e-value cutoff 0.1) to search for the neighbor genes longest isoform tophit in the *C. elegans* proteome. We then found what chromosome the *C. elegans* tophit was coded on. These neighbor gene *C. elegans* tophit chromosome is what is plotted for each confident GATA-domain-containing gene in Figure 4.

### Testing for changes in selection pressure intensity

RELAX (Wertheim et al. 2015) compares two sets of designated branches in a phylogeny and tests whether the data is better fit by a single distribution of three non-synonymous substitutions per site to the number of synonymous substitutions per site (dN/dS) rate categories among all branches or by different distributions for each set where the rate categories in one are related to the rate categories in the other by an exponentiation factor (k). We used RELAX to evaluate selection intensity changes in the rcor1 and elt6 clades because they our results suggests both of these clades expanded through gene duplication in the Caenorhabditis genus. There were a few species-specific paralogs where one protein was more conserved and the other(s) more divergent (black bars in Figure 5). In these cases, we only included the more conserved one, which made our tests more conservative. We used three possible rate categories in each test and the default settings.

### Data availability

Custom Python and R scripts used for this article will be shared on reasonable request to the corresponding author.

## Acknowledgements

Thanks to The Caenorhabditis Genome Project for *Caenorhabditis* and outgroup datasets. Special thanks to Lewis Stevens for pre-publication access to *Caenorhabditis* genomic data. We gratefully acknowledge Rifkin lab members for their insightful discussions. This work was supported by the National Institutes of Health (R01 GM103782); and the National Science Foundation (IOS 1936674).

## Abbreviations

DBD: DNA-binding domains
GATA factors: GATA-type transcription factors
SANT: Swi3/Ada2/N-CoR/TFIIIB
ELM2: EGL-27 and MTA1 homology 2
ZnF: zinc finger
pHMM: profile hidden Markov model
cGATA-1: chicken GATA-1
CF: C-terminal zinc finger
hGATA-3: human GATA-3
BAH: bromo-adjacent homology
bp: base pair
BR: basic region
NF: N-terminal zinc finger
ZnF tree: GATA-domain-only phylogeny
NJ: neighbor-joining
dN/dS: ratio of the number of non-synonymous substitutions per site to the number of synonymous substitutions per site

---

**Supplemental Table 1. Additional information on 941 GATA-domain-containing proteins and the genes encoding them.** (See separate large csv file).

**Supplemental Table 2. Number of GATA-domain-containing proteins in each species.** (See separate large csv file).

**Supplemental Table 3. Syntenic GATA-domain-containing proteins.** (See separate large csv file).

**Supplemental Figure 1.**
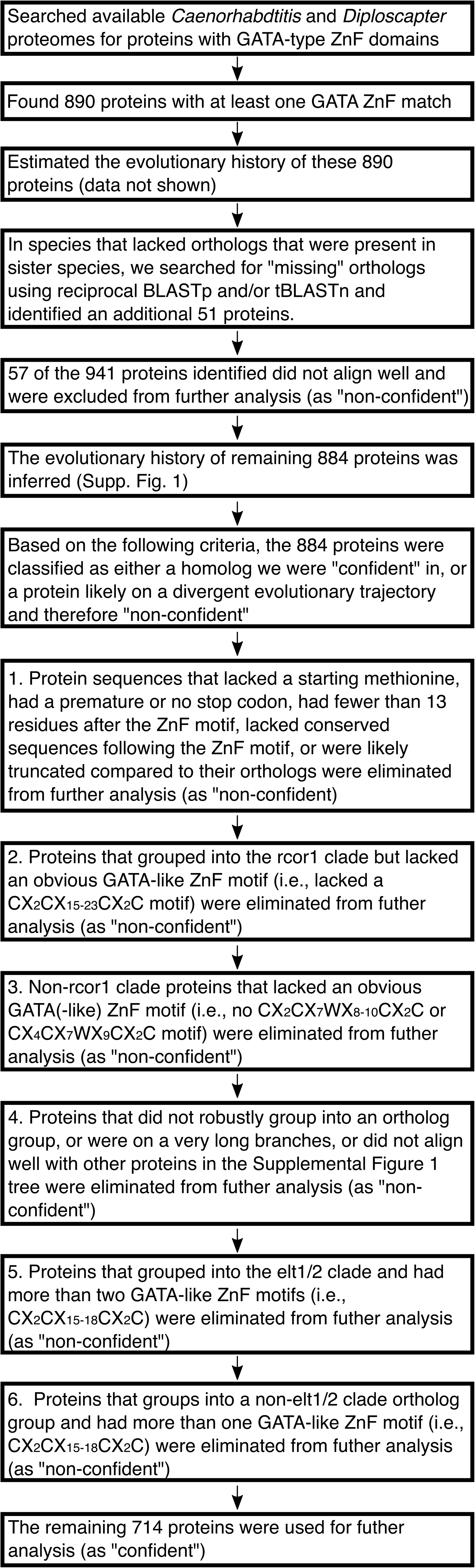
Method of selecting protein sequences for further analyses. The process carried out to select among putative GATA-domain-containing protein sequences those for further analyses are depicted in this figure. Protein sequences were classified as “confident” for use in further analyses versus “not_confident” using this decision tree and its selection criteria. (Supplemental Table 1 comprises a list of the resulting classifications for each protein sequence.) ZnF stands for zinc finger.

**Supplemental Figure 2.**
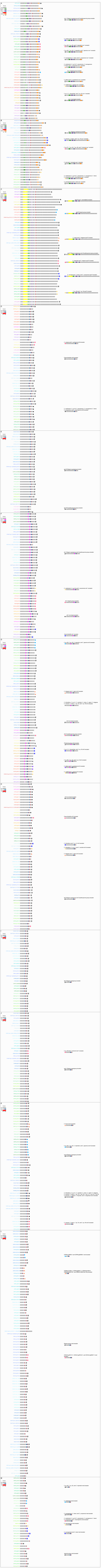
Comparison of extant gene structures and predictions of the structures of their ancestral genes. Gene structures for proteins in this study deemed “confident” (see Materials and Methods and Supplemental Table 1) are shown on the left side of each panel. The species from which each gene sequence was obtained is indicated by the font color of the name given to each sequence (which appears to the left of each gene structure); color-species designations are the same as those used in Fig. 1B). A key to the color coding of the protein domains depicted in the gene structures is shown at the top left of each panel. The genes in each panel are organized in the order that they appear in our phylogeny (Fig. 1A), i.e., in relation to their relatedness to each other. Predicted ancestral gene structures are shown on the right side of each panel); their names (depicted above each ancestral gene structure) reflect the species their predicted structures were based on. The genes used to calculate median exon lengths in the predicted ancestral gene structures are identified with matching symbols (placed at the end of the former and the start of the latter). Structures of extant genes that have a divergent number of exons from the predicted ancestral gene structures do not have a symbol next to them. The structures of all the genes comprising a single ortholog group are depicted in separate, individual panels, with the exception of the MED ortholog group; the *meds* structures are depicted in two panels (D and E), one containing the structures of *meds* genes from species in the *Elegans* group and the other those from species in the *Japonica* group. The gene family represented in each panel is as follows: **(A)** *rcor-1*; **(B)** *spr-1*; **(C)** *egl-27*; **(D)** *elt-6*; **(E)** *egl-18*; **(F)** *elt-1*; **(G)** *elt-2*; **(H)** *elt-3*; **(I)** *egl-18*; **(J)** *end-1*; **(K)** *end-3*; **(L)** *Elegans meds*; and **(M)** *Japonica meds*.

**Supplemental Figure 3.**
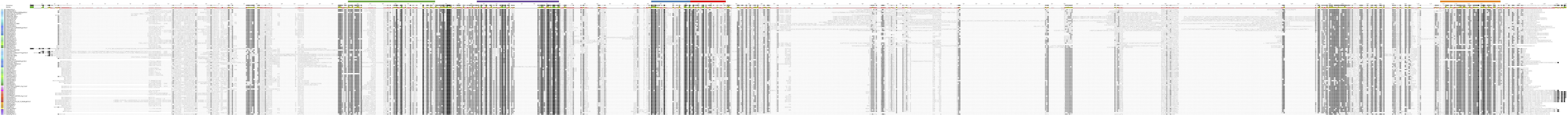
Full-length alignments of rcor1 clade proteins. Muscle-aligned protein sequences for confident rcor1 clade proteins are shown in the order of the Stevens 2020 species phylogeny. Residues are shaded by similarity. Each consensus sequence is shown at the top of each panel. The percent identity to the consensus is plotted underneath the consensus sequence. Protein domains encoded by the sequences are highlighted above the alignment as colored rectangles. The domain colors are the same as used in Figure 2 and Supplemental Figure 2 (i.e., ZnF in blue, BR in red, ELM2 in green, Myb1/SANT1 in purple, and Myb2/SANT2 in orange). The gene names are shown on the left of the alignment. A vertical bar to the left of the gene names is colored by ortholog group. The ortholog group color-coding is the same as in Figure 4 (i.e., SPR-1s with dark gray and RCOR-1s with light gray). The species colors are shown to the left of the ortholog group bar (which are the same as in Figure 1B).

**Supplemental Figure 4.**
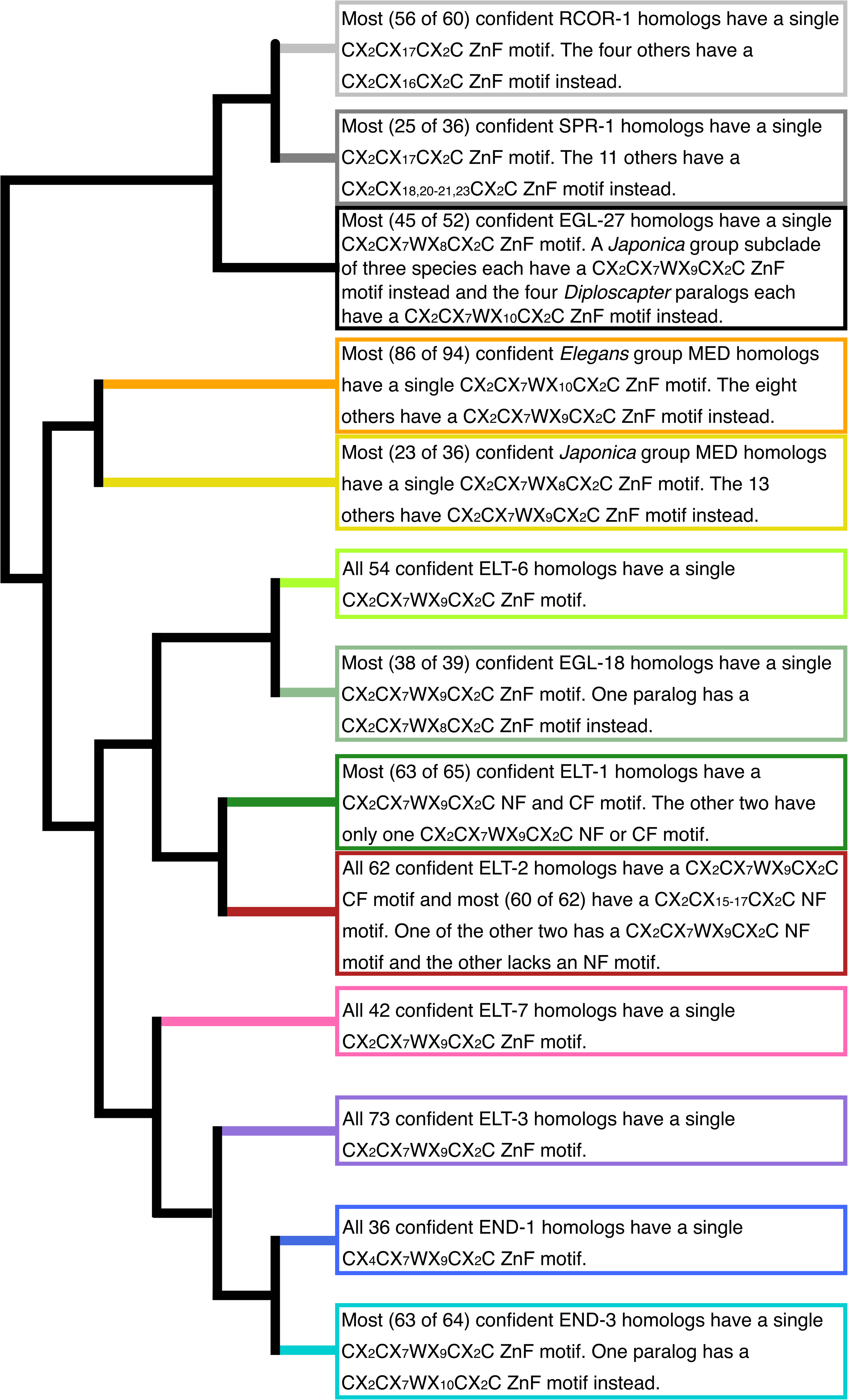
Comparison of zinc finger motifs found in each ortholog group. The loop size(s) of each GATA factor ZnF, and the presence or absence of the highly conserved tryptophan (W) at position eight in the ZnF loop, is shown for all genes deemed confident in each of the 12 ortholog groups.

**Supplemental Figure 5.**
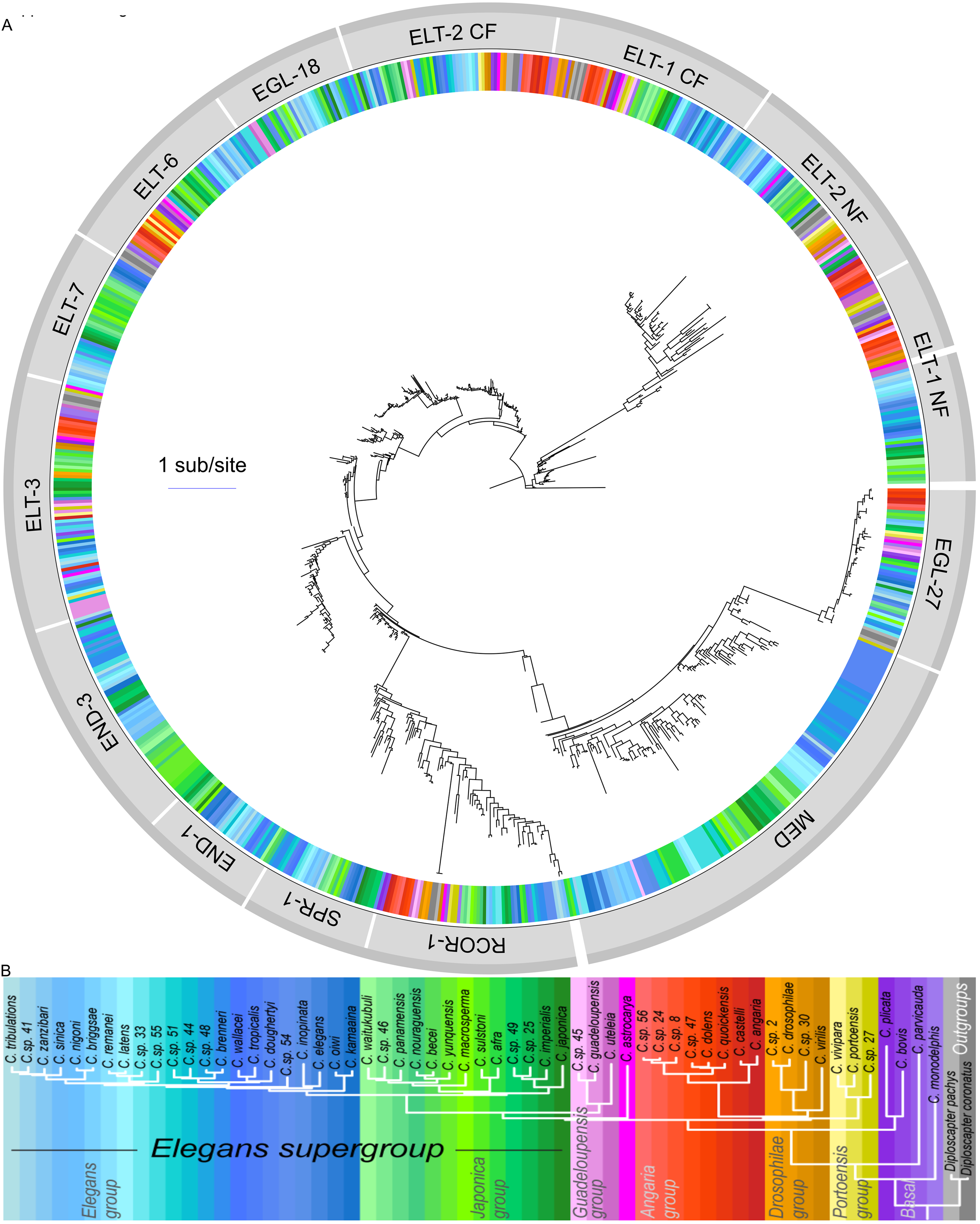
Phylogenetic analysis of GATA (or GATA-like) domains. Maximum likelihood phylogeny of confident GATA domains in 58 *Caenorhabditis* and two outgroup nematode species. The GATA domain from a *D. fasciculatum* (slime mold) GATA factor was used to root the tree. The colors in the ring encircling the tree correspond to the species in which the GATA domain is from (the key to color-species correspondence is the same as in Supplemental Figure 1B). The 14 different groups that the GATA domain cluster into are labeled in light gray bars on the outside of the species color ring. Groupings of ortholog groups that share adjacent clades in the tree are highlighted by a darker gray line on the outermost edge of the figure. The key for translating branch length into evolutionary distance (in units of amino acid substitutions per site) is shown near the bottom of the phylogenetic tree.

**Supplemental Figure 6.**
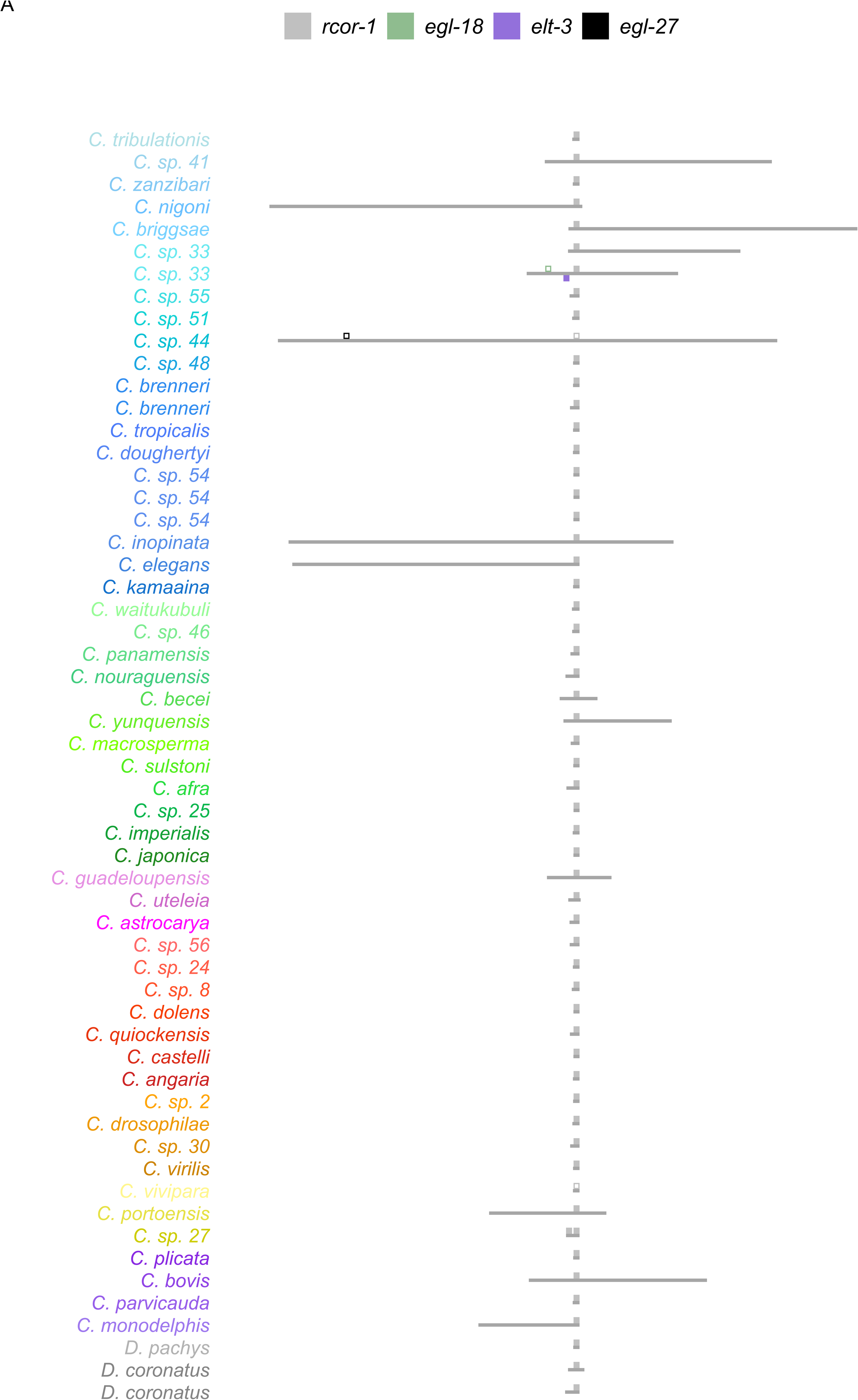
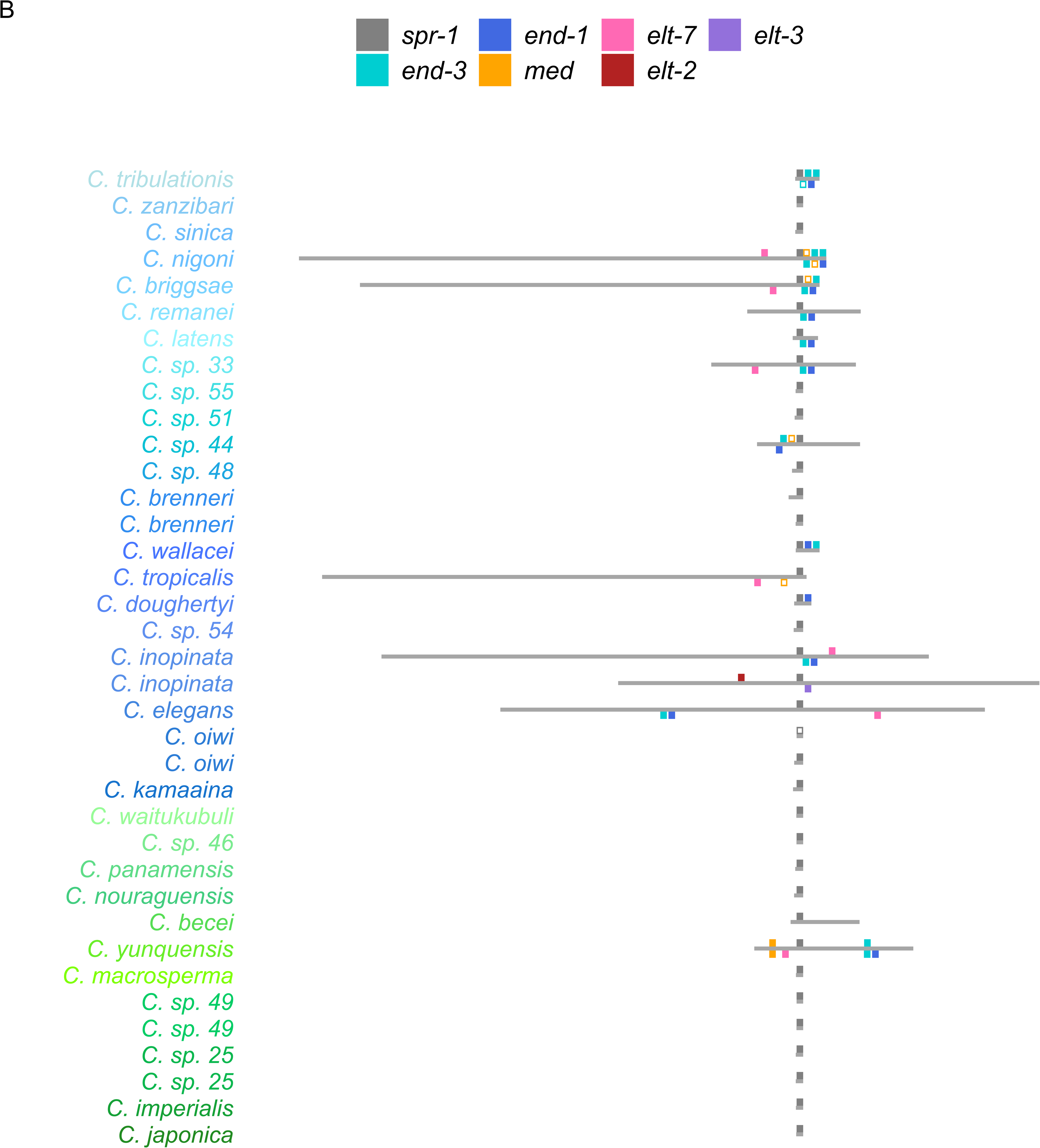
Contig/scaffold/chromosome locations of rcor1 clade genes. Contigs/scaffolds/chromosomes (depicted as gray horizontal rectangles) are anchored on a respective GATA-domain-containing gene (depicted as colored squares). The relative locations of any other GATA- domain-containing genes (depicted as other color squares) on the same scaffold/chromosome (i.e., syntenic GATA-domain-containing genes) are shown above or below a given contig/scaffold/chromosome, indicating their orientation on the same or opposite strand, respectively, as the anchored gene. Genes deemed confident and non-confident are depicted as filled in or outlined colored squares, respectively. Genes from each ortholog group are designated using the same color, as noted in the key at the top of each plot. The species from which each respective contig/scaffold/chromosome was sequenced is indicated on its left. The species names are in the order of the species phylogeny (Stevens 2020) and color-coded as in Figure 1B. (For visual clarity, the sizes and exact relative locations of the colored squares representing GATA-domain-containing genes have been adjusted slightly in some cases, and large contigs/scaffolds/chromosomes were scaled down (based on their actual length per plot) while the smallest contigs/scaffolds were lengthened.) The gene serving as the anchor in each panel is as follows: **(A)** *rcor-1* and **(B)** *spr-1*.

**Supplemental Figure 7.**
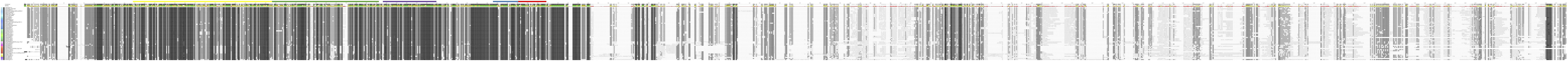
Full-length alignments of EGL-27 proteins. Muscle-aligned protein sequences for confident EGL-27 ortholog group proteins are shown in the order of the Stevens 2020 species phylogeny. Residues are shaded by similarity. Each consensus sequence is shown at the top of each panel. The percent identity to the consensus is plotted underneath the consensus sequence. Protein domains encoded by the sequences are highlighted above the alignment as colored rectangles. The domain colors are the same as used in Figure 2 and Supplemental Figure 2 (i.e., ZnF in blue, BR in red, BAH in yellow, ELM2 in green, and Myb1/SANT1 in purple). The gene names are shown on the left of the alignment. The species colors are shown to the left of the ortholog group bar (which are the same as in Figure 1B).

**Supplemental Figure 8.**
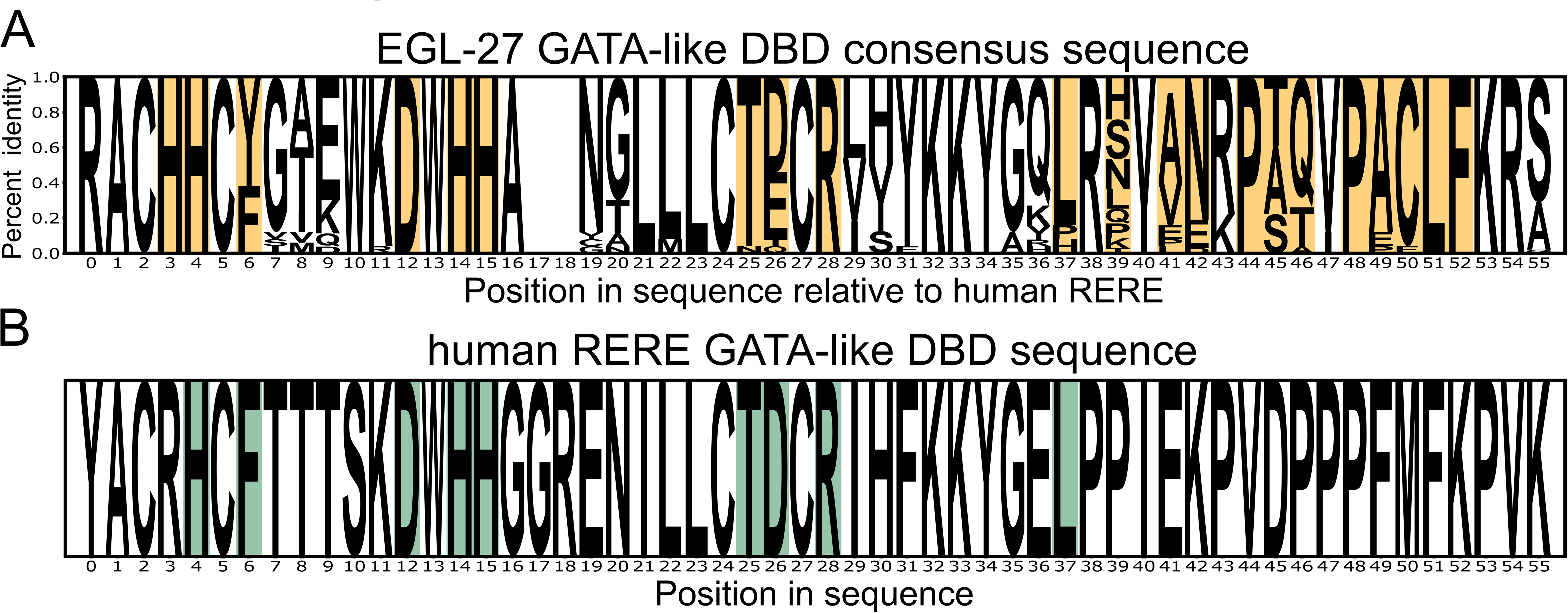
Comparison of GATA DBD sequences in *Caenorhabditis* EGL-27 and human RERE. **(A)** The consensus GATA DBD for *Caenorhabditis* EGL-27. Ten residues not found in any other GATA DBDs in *Caenorhabditis* species are highlighted in light orange. (Gaps were included at positions 17 and 18 to provide alignment with the human RERE protein.) **(B)** The GATA DBD sequence of human RERE. The nine (of the 10) residues specific to the consensus *Caenorhabditis* EGL-27 DBD are highlighted in light blue.

**Supplemental Figure 9.**
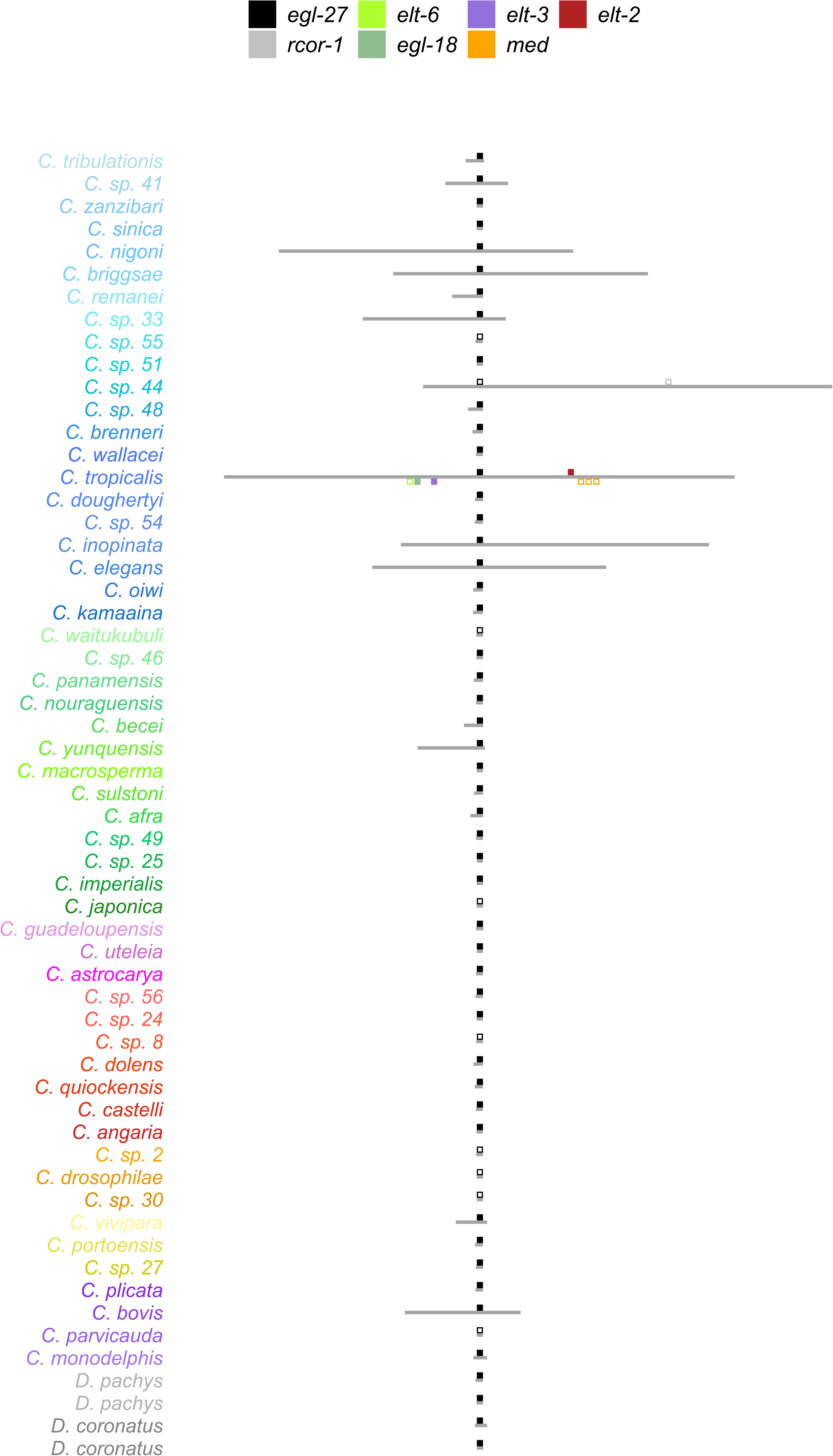
Contig/scaffold/chromosome locations of EGL-27 ortholog group genes. Contigs/scaffolds/chromosomes (depicted as gray horizontal rectangles) are anchored on a respective GATA-domain-containing gene (depicted as colored squares). The relative locations of any other GATA- domain-containing genes (depicted as other color squares) on the same scaffold/chromosome (i.e., syntenic GATA-domain-containing genes) are shown above or below a given contig/scaffold/chromosome, indicating their orientation on the same or opposite strand, respectively, as the anchored gene. Genes deemed confident and non-confident (see Materials and Methods) are depicted as filled in or outlined colored squares, respectively. Genes from each ortholog group are designated using the same color, as noted in the key at the top of each plot. The species from which each respective contig/scaffold/chromosome was sequenced is indicated on its left. The species names are in the order of the species phylogeny (Stevens 2020) and color-coded as in Figure 1B. (For visual clarity, the sizes and exact relative locations of the colored squares representing GATA-domain-containing genes have been adjusted slightly in some cases, and large contigs/scaffolds/chromosomes were scaled down (based on their actual length per plot) while the smallest contigs/scaffolds were lengthened.)

**Supplemental Figure 10.**
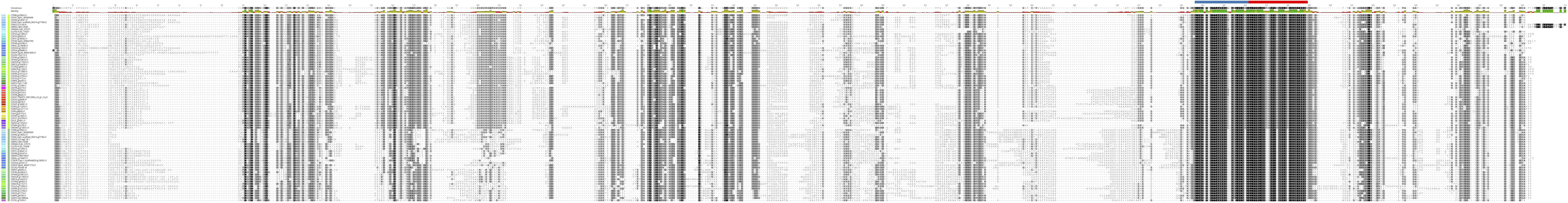
Full-length alignments of elt6 clade proteins. Muscle-aligned protein sequences for confident elt6 clade proteins are shown in the order of the Stevens 2020 species phylogeny. Residues are shaded by similarity. Each consensus sequence is shown at the top of each panel. The percent identity to the consensus is plotted underneath the consensus sequence. Protein domains encoded by the sequences are highlighted above the alignment as colored rectangles. The domain colors are the same as used in Figure 2 and Supplemental Figure 2 (i.e., ZnF in blue and BR in red). The gene names are shown on the left of the alignment. A vertical bar to the left of the gene names is colored by ortholog group. The ortholog group color-coding is the same as in Figure 4 (i.e., ELT-6s with lime green and EGL-18s with sea green). The species colors are shown to the left of the ortholog group bar (which are the same as in Figure 1B).

**Supplemental Figure 11.**
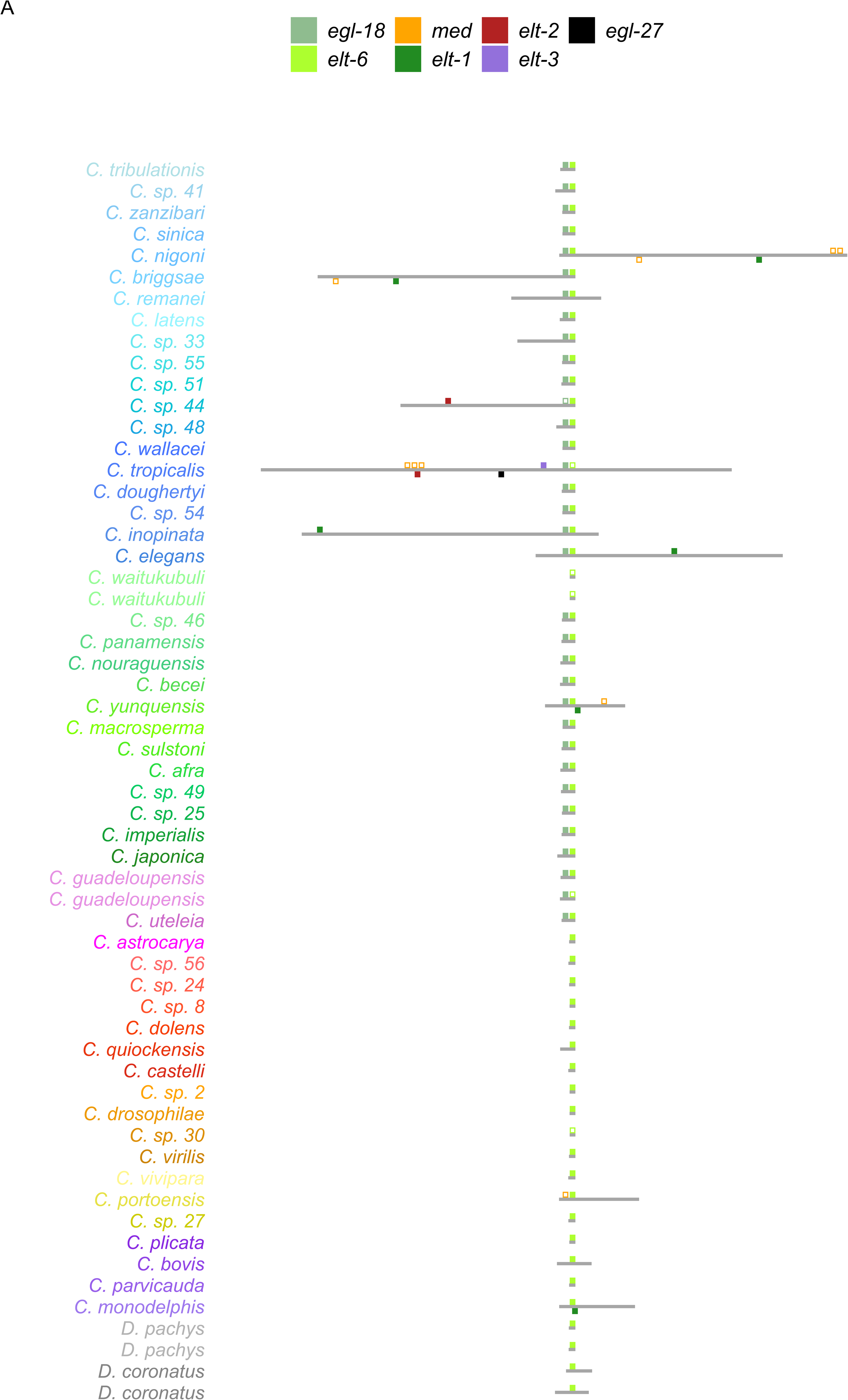
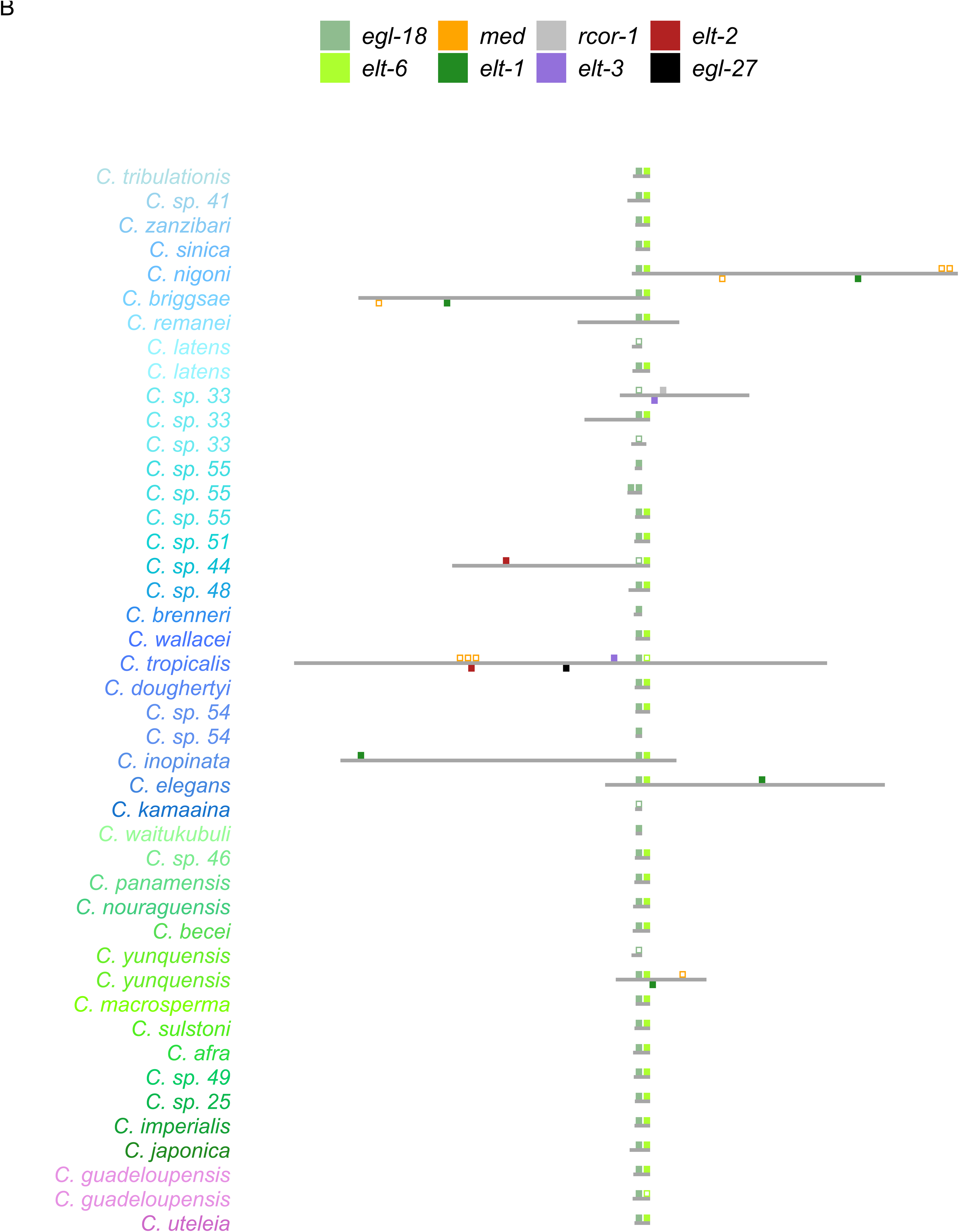
Contig/scaffold/chromosome locations of elt6 clade genes. Contigs/scaffolds/chromosomes (depicted as gray horizontal rectangles) are anchored on a respective GATA-domain-containing gene (depicted as colored squares). The relative locations of any other GATA- domain-containing genes (depicted as other color squares) on the same scaffold/chromosome (i.e., syntenic GATA-domain-containing genes) are shown above or below a given contig/scaffold/chromosome, indicating their orientation on the same or opposite strand, respectively, as the anchored gene. Genes deemed confident and non-confident (see Materials and Methods) are depicted as filled in or outlined colored squares, respectively. Genes from each ortholog group are designated using the same color, as noted in the key at the top of each plot. The species from which each respective contig/scaffold/chromosome was sequenced is indicated on its left. The species names are in the order of the species phylogeny (Stevens 2020) and color-coded as in Figure 1B. (For visual clarity, the sizes and exact relative locations of the colored squares representing GATA-domain-containing genes have been adjusted slightly in some cases, and large contigs/scaffolds/chromosomes were scaled down (based on their actual length per plot) while the smallest contigs/scaffolds were lengthened.) The gene serving as the anchor in each panel is as follows: **(A)** *elt-6*; and **(B)** *egl-18*.

**Supplemental Figure 12.**
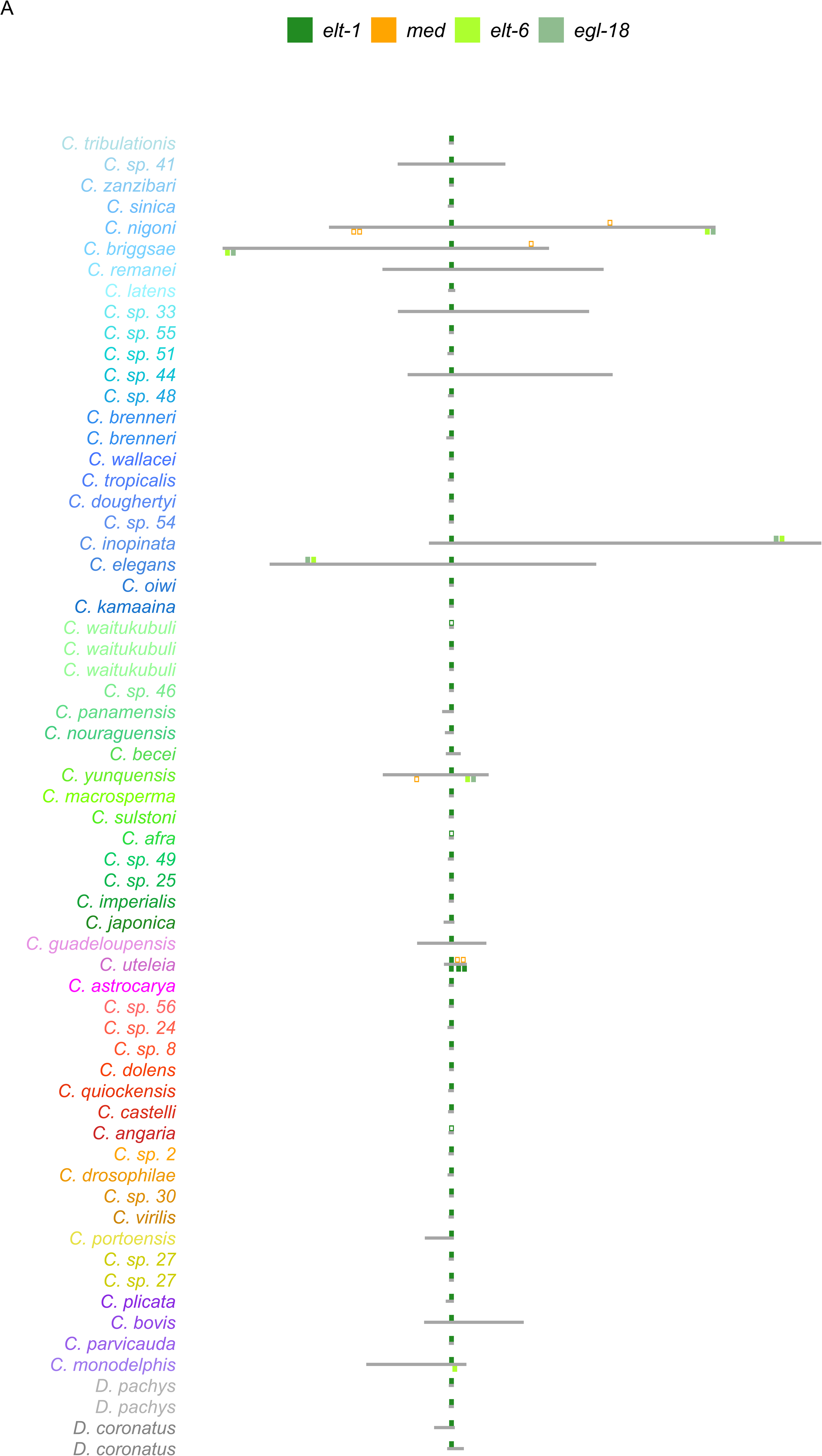
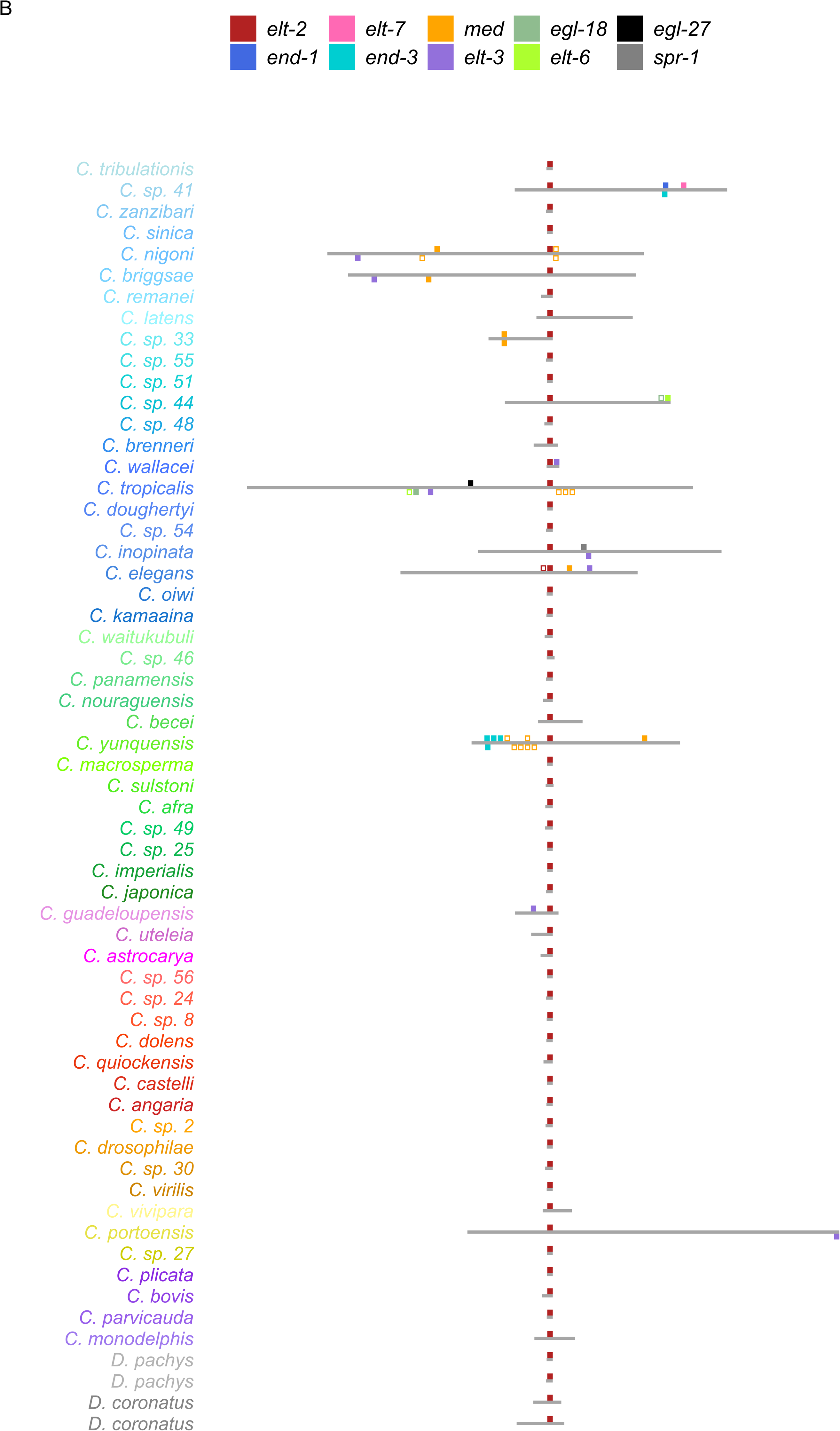
Contig/scaffold/chromosome locations of elt1/2 clade genes. Contigs/scaffolds/chromosomes (depicted as gray horizontal rectangles) are anchored on a respective GATA-domain-containing gene (depicted as colored squares). The relative locations of any other GATA- domain-containing genes (depicted as other color squares) on the same scaffold/chromosome (i.e., syntenic GATA-domain-containing genes) are shown above or below a given contig/scaffold/chromosome, indicating their orientation on the same or opposite strand, respectively, as the anchored gene. Genes deemed confident and non-confident (see Materials and Methods) are depicted as filled in or outlined colored squares, respectively. Genes from each ortholog group are designated using the same color, as noted in the key at the top of each plot. The species from which each respective contig/scaffold/chromosome was sequenced is indicated on its left. The species names are in the order of the species phylogeny (Stevens 2020) and color-coded as in Figure 1B. (For visual clarity, the sizes and exact relative locations of the colored squares representing GATA-domain-containing genes have been adjusted slightly in some cases, and large contigs/scaffolds/chromosomes were scaled down (based on their actual length per plot) while the smallest contigs/scaffolds were lengthened.) The gene serving as the anchor in each panel is as follows: **(A)** *elt-1*; and **(B)** *elt-2*.

**Supplemental Figure 13.**
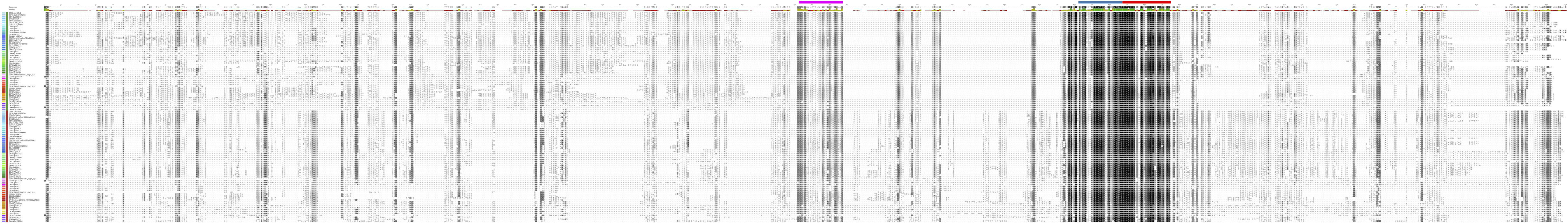
Full-length alignments of elt1/2 clade proteins. Muscle-aligned protein sequences for confident elt1/2 clade proteins are shown in the order of the Stevens 2020 species phylogeny. Residues are shaded by similarity. Each consensus sequence is shown at the top of each panel. The percent identity to the consensus is plotted underneath the consensus sequence. Protein domains encoded by the sequences are highlighted above the alignment as colored rectangles. The domain colors are the same as used in Figure 2 and Supplemental Figure 2 (i.e., CF/ZnF in blue, BR in red, and NF in pink). The gene names are shown on the left of the alignment. A vertical bar to the left of the gene names is colored by ortholog group. The ortholog group color-coding is the same as in Figure 4 (i.e., ELT-1s with green and ELT-2s with red). The species colors are shown to the left of the ortholog group bar (which are the same as in Figure 1B).

**Supplemental Figure 14.**
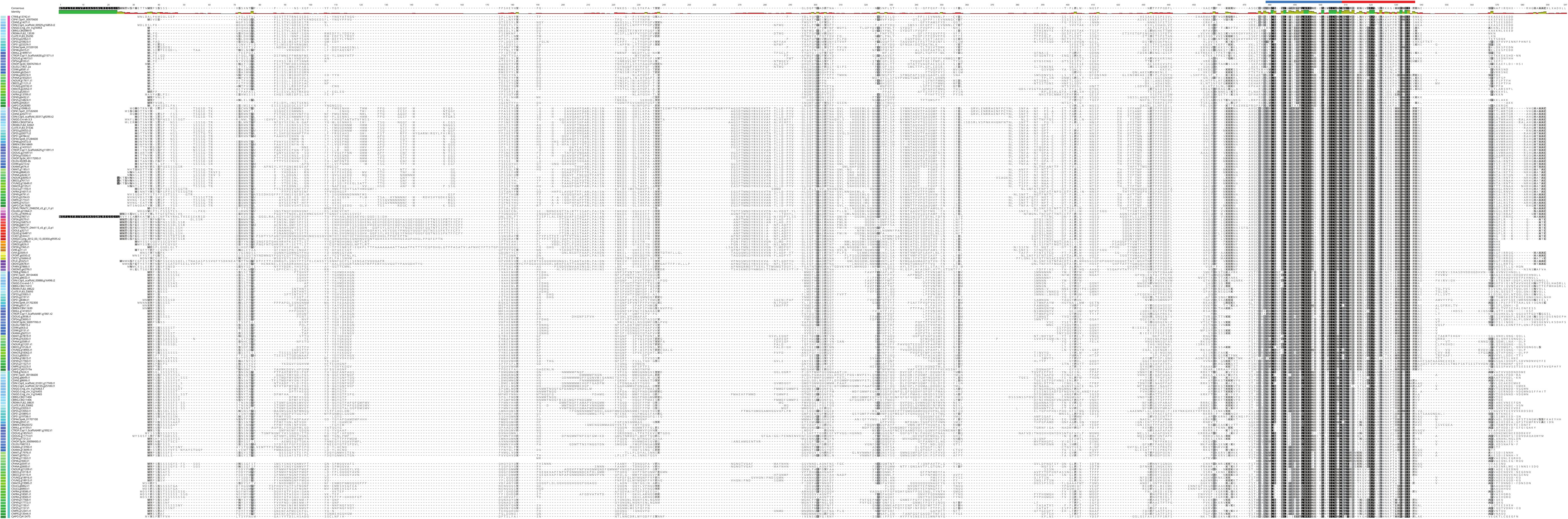
Full-length alignments of elt3 clade proteins. Muscle-aligned protein sequences for confident representative elt3 clade proteins are shown in the order of the Stevens 2020 species phylogeny. Residues are shaded by similarity. Each consensus sequence is shown at the top of each panel. The percent identity to the consensus is plotted underneath the consensus sequence. Protein domains encoded by the sequences are highlighted above the alignment as colored rectangles. The domain colors are the same as used in Figure 2 and Supplemental Figure 2 (i.e., ZnF in blue and BR in red). The gene names are shown on the left of the alignment. A vertical bar to the left of the gene names is colored by ortholog group. The ortholog group color-coding is the same as in Figure 4 (i.e., ELT-7s with pink, ELT-3s with purple, END-1s with blue, and END-3s with teal). The species colors are shown to the left of the ortholog group bar (which are the same as in Figure 1B).

**Supplemental Figure 15.**
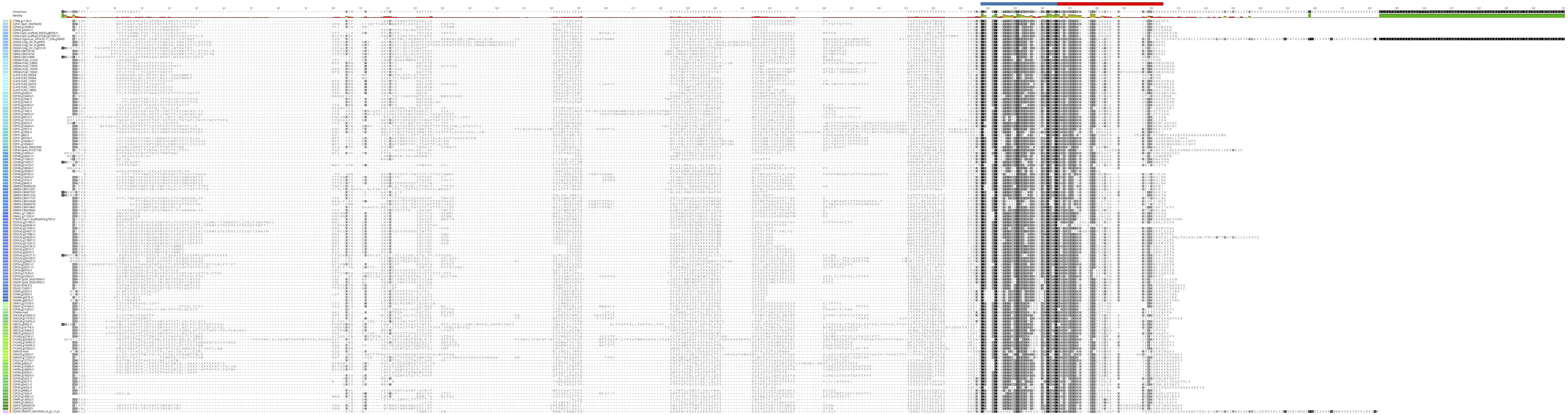
Full-length alignments of MED proteins. Muscle-aligned protein sequences for confident MED ortholog group proteins are shown in the order of the Stevens 2020 species phylogeny. Residues are shaded by similarity. Each consensus sequence is shown at the top of each panel. The percent identity to the consensus is plotted underneath the consensus sequence. Protein domains encoded by the sequences are highlighted above the alignment as colored rectangles. The domain colors are the same as used in Figure 2 and Supplemental Figure 2 (i.e., ZnF in blue and BR in red). The gene names are shown on the left of the alignment. The species colors are shown to the left of the ortholog group bar (which are the same as in Figure 1B).

**Supplemental Figure 16.**
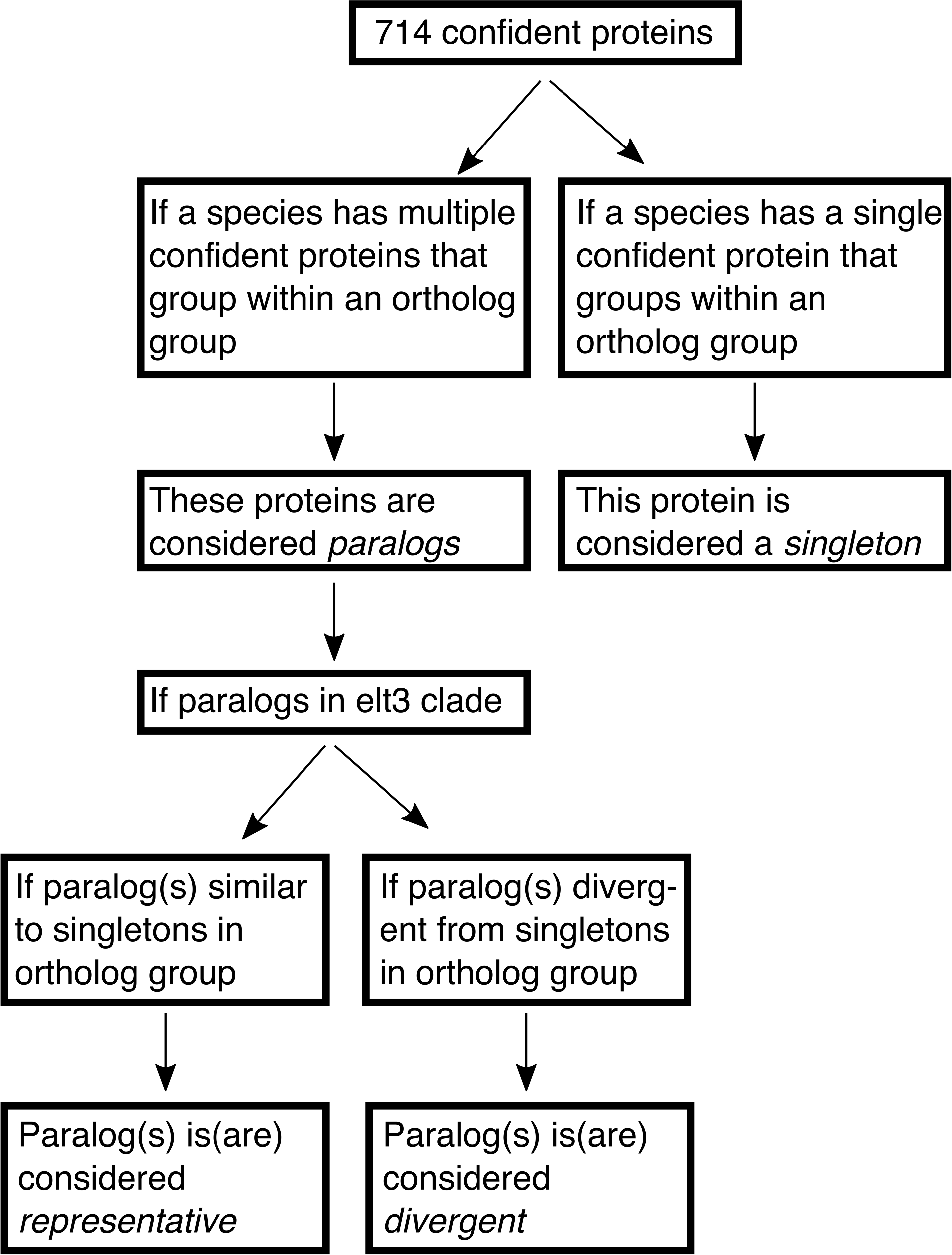
Classification of paralogs in the Elt3 clade. Decision tree used for classifying paralogs in the elt3 clade as representative or divergent.

